# A New anticancer substance from cancer cell apoptosis. CSS (Cancer cell Suicide Substance) production in culture

**DOI:** 10.1101/2025.10.20.683431

**Authors:** Tomoyuki Tajima, Yoshifusa Kondo

**Affiliations:** Medical Corporation Ichikawa Clinic-Affiliated Biomedical Research Institute, Medical Corporation Ichikawa Clinic, 1-17-19 Hirata, Ichikawa city, Chiba 272- 0031, Japan

**Keywords:** cancer, cancer cell death, anticancer, apoptosis, intercellular communication, cancer therapy

## Abstract

It is well-known that the cells die when a cancer cell line is continuously cultured without passage, even though it was kept in a fresh medium. However, none of this phenomenon’s decisive causes and mechanism(s) have been elucidated yet. After the cancer cell death, the medium exserted cytocidality. This cancer cell death (e.g., spontaneous, apoptosis, programmed) was found to be caused by only one kind of cytocidal substance (MW<1kDa) (named CSS) produced from the cells own component(s) by the cells themselves. CSS mammalian specifically by cancer cell lines but not nontumor cell lines and the CSS exerted cytocidality against other cell lines and vice versa regardless of species in mammalian. In vivo, CSS showed a significant life-prolonging effect without any adverse event. An established cell line derived from <u>h</u>uman renal cell <u>c</u>arcinoma (HRC23) was used in this study as CSS was found and characterized by this cell line first.

**Highlights:**

1. Only one kind of low molecular substance exerting cytocidality was found only in a medium after cancer cells death in the culture but not in non-tumor cell lines one.
2. The cytocidal substance (named CSS: **<u>c</u>**ancer cell **<u>s</u>**uicide **<u>s</u>**ubstance) was a small molecule (MW<1kDa) with a slightly positively charged highly hydrophilic character.
3. CSS was produced regardless of extracellular nutritional condition, even in glucose-free HBSS (**Nutrition-Free Method.’**)
4. CSS exserted cytocidality against other cancer cell lines irrespective of the histological patterns and the species of mammalian cells and vice versa.
5. CSS can be considered as a direct and critical substance produced from cell component(s) themselves by the cancer cells themselves.
6. Production of CSS was specific for cancer cells. This clear functional difference between cancer and nontumor cells should be meaningful embryologically.
7. ***In vivo***, CSS showed complete anti-metastasis during administration period against <u>L</u>ewis <u>L</u>ung <u>C</u>arcinoma (LLC) transplanted mouse without any adverse event. CSS may have high potential as a new type of cancer therapy drug.

**In short:** This study is so simple, absolutely reproducible and anyone can reproduce this study with an easy basic method. A critical substance CSS (<u>c</u>ancer cell <u>s</u>uicide <u>s</u>ubstance) of spontaneous cancer cell death was found in a medium after the cancer cells had been dead. Only one kind of small-size substance with MW<1kDa with a highly water-soluble and positively charged character is made only from cell own component(s) itself by the cells themselves. Even in a nutrition-free physiologically balanced salt solution, CSS was produced regardless of extracellular nutritional condition. Only cancer cell lines produced CSS, but not by non-tumor cell lines. CSS exserted cytocidality against other histologically different patterns of cancer cell lines regardless of species and vice versa. The anticancer effect of HRC23 produced CSS in vivo against mouse LLC (Lewis Lung Carcinoma)-transplanted mice showed a clear life-prolonging effect without any adverse event. Thus, CSS may have the possibility to be a cancer therapy drug. And, in view of embryologically, this clear difference of CSS production between cancer cells and nontumor cells will be important event.

## Introduction

This study was based on why cultured cancer cell lines of established lines require passage to maintain an infinite life span. No definitive cause or mechanisms for this spontaneous cancer cell death (this cell death is called *“****terminal death****” i*n this study) have not been elucidated yet.^1, 2, 3, 13, 14^ Little attention seems to have been paid to this phenomenon as far as authors know. When a culture reaches the confluent phase, the cells become overpopulated by continuous cell growth. This phase is called the “terminal phase” here. Further cultivation with frequent medium changes to fresh medium without passage, cell death inevitably occurs. This death is called *“****terminal death****”* in this study. The deterioration of nutrition in the culture environment, which has been commonly considered, seemed neither to be one nor only cause of *“****terminal death***.*”* Therefore, the relationship between the nutritional environment (as an extracellular factor) and *“****terminal death****”* in culture was investigated. This study used an established cell line derived from <u>h</u>uman renal cell <u>c</u>arcinoma (HRC23; the same as ku-2 cell line ^**5, 6, 7**^.)

## Results

Assuming some kinds of substances cause the “***terminal death***” is produced. The following experiments were performed to investigate the existence of cytocidal substance(s).

### 1. Examination of nutritional conditions on the cytocidal effect of media after “*terminal death*” using three kinds of solutions for the ‘final solution replacement’ at the “terminal phase” in HRC23 culture (Figures. 1: A. and B.) ^17.17.19. 20^

**“*Terminal death”*** was recognized when HRC23 was cultured only by medium exchange without passage. After the cell death (**“*terminal death***,**”**) the medium was taken from the cultured bottle, and nutrients and serum were added to the supernatant of collected cell-dead medium with pH adjustment (‘regeneration’ procedure for reuse as culture medium.) The cells died when the fresh culture was exchanged with this regenerated medium. Moreover, the cytocidal effect showed concentration dependency. Hence, the existence of some cytocidal component was suggested. **(Figures 2 and Figure 3: EMEM + FBS)** Then the relationship between the nutritional environment as an extracellular condition and **“*terminal death*”** was examined.

**Figure 1.**
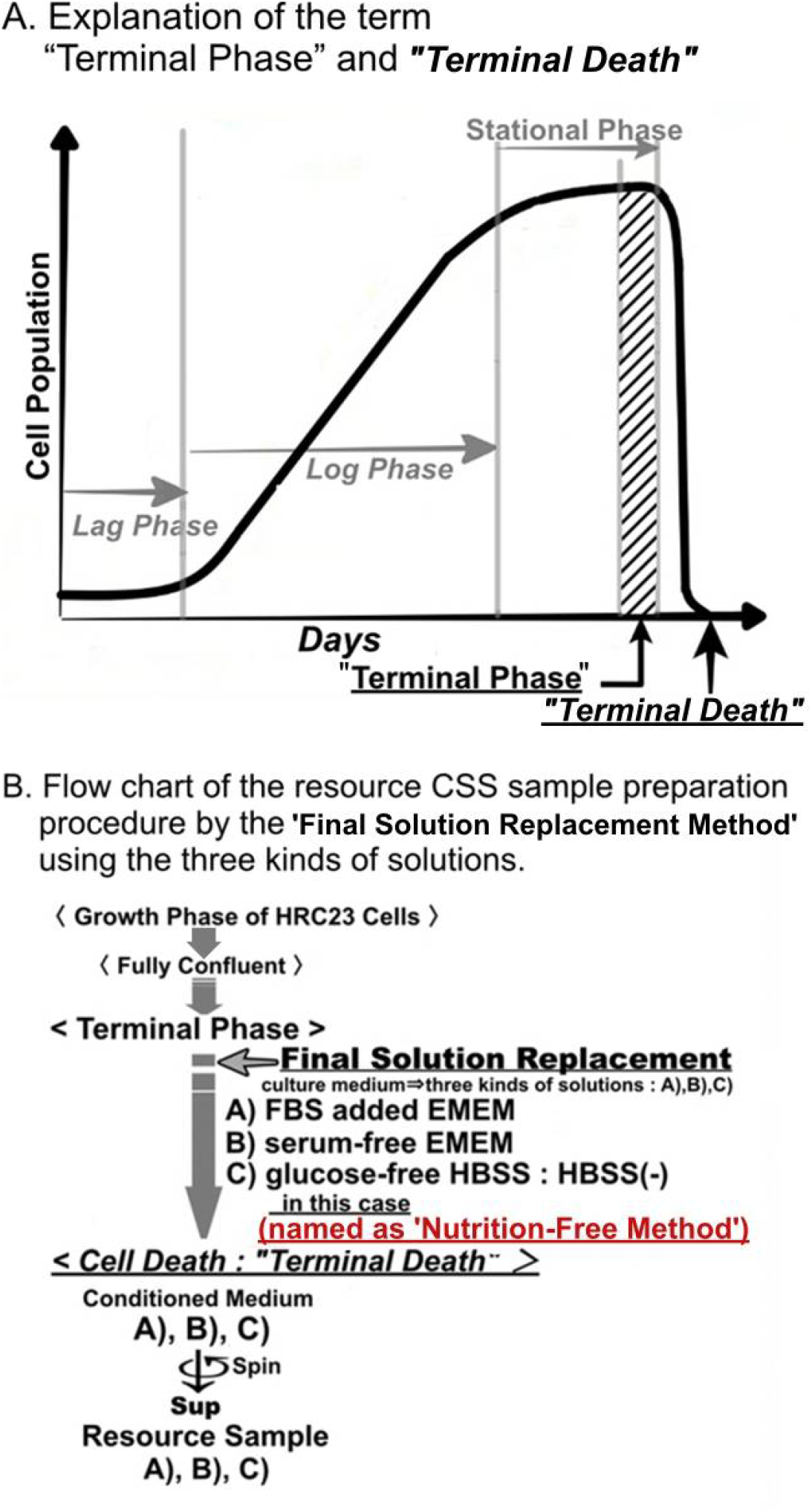
A: Explanation of the term *“*Terminal Phase*”* and *“Terminal Death”* in a cell culture. B: Flow chart of the sample preparation procedure by final solution replacement using the three kinds of solutions. **Solutions: A: EMEM+FBS, B: FBS-free EMEM, and C: HBSS (-).** The medium of HRC23 cells cultured until fully confluent state (“*terminal phase*”) was replaced with the 3 kinds of solutions as the final replacement solution, respectively. Then the cells in each solution were further cultured until they died (**“*terminal death*.”**) The cells-dead each solution was aseptically collected, and their supernatant was used as the sample to measure the cytocidal effect. For the sample made with serum-free EMEM and HBSS (-), half the volume of each sample was fractionated by ultrafiltration using a molecular weight cut membrane (nominal molecular cut weight is 1 kDa) to examine the molecular size of cytocidal effect exserting component(s.) The result is shown in (**Figure 3**.)

**Figure 2.**
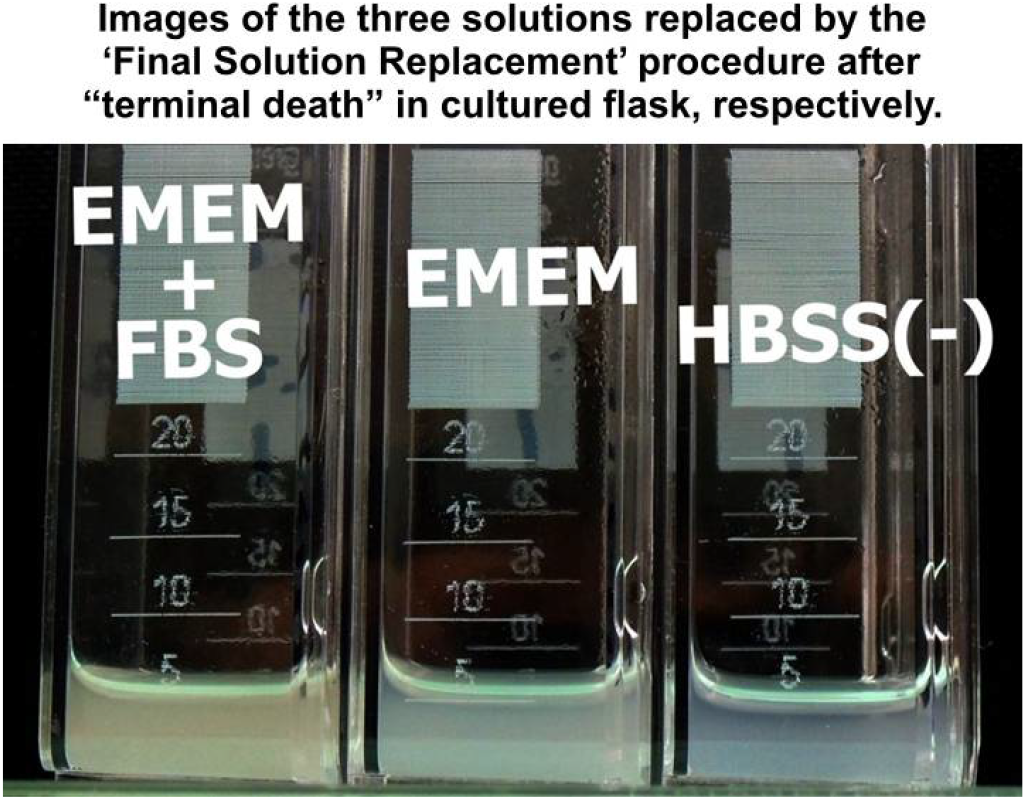
Images of the three solutions replaced by the final solution replacement procedure after *“terminal death”* in cultured flask, respectively. left: EMEM+FBS, center: EMEM, right: HBSS (-) · (see also **Figure. 1. B**) Cells were detached from the adhesive surface of the culture flask, and the three solutions were turbid with cell fragments.

**Figure 3.**
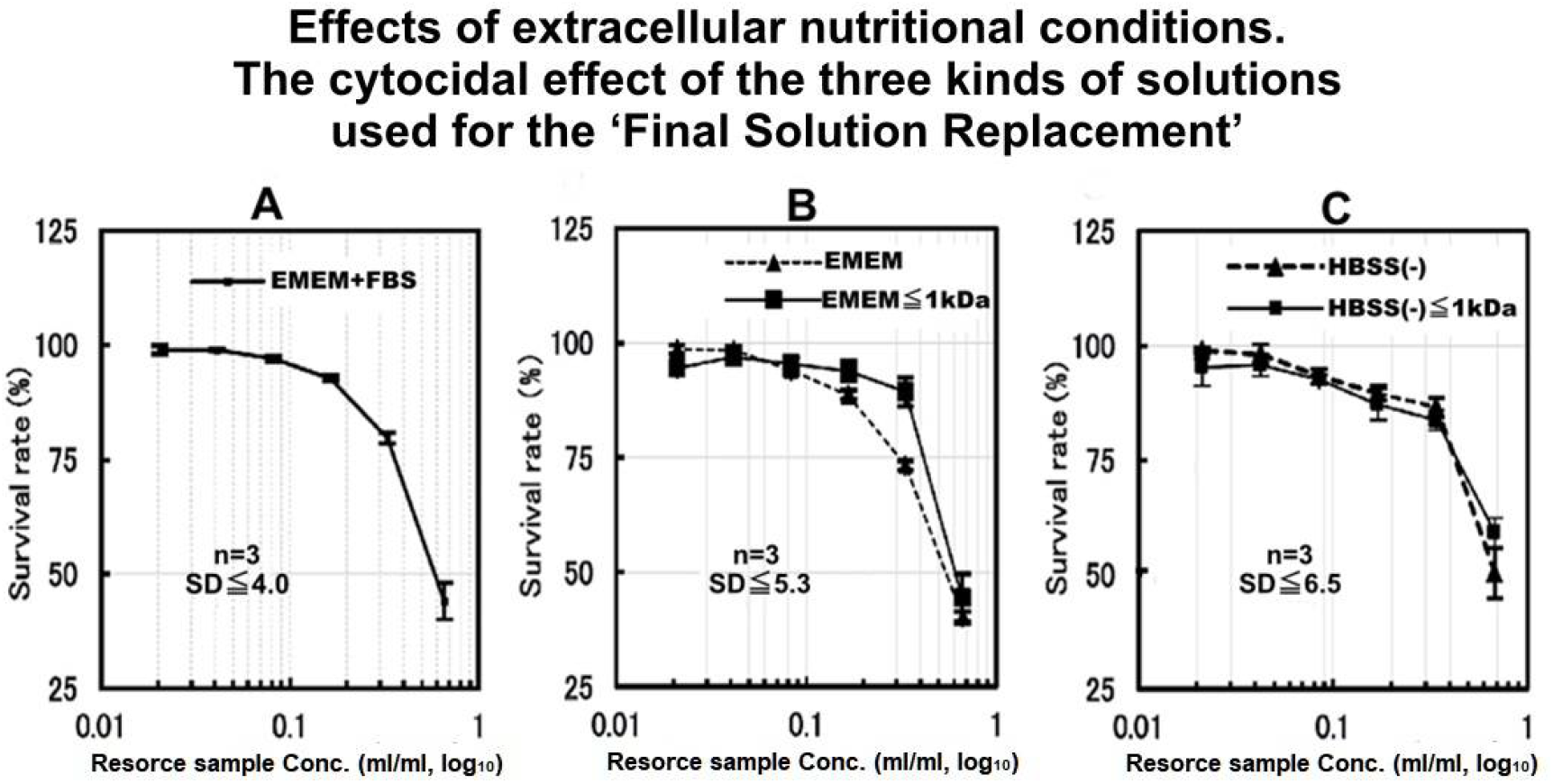
The nutritional requirement for CSS production of the cytocidal effect of different solutions obtained after cell death was investigated using the final solution replacement method using the three kinds of solutions by an MTT assay. Cytocidal effect of the three kinds of collected and regenerated solutions against fresh HRC23 culture, those were used at final replacement **[A: 10%FBS-added EMEM, B: EMEM only, and C: HBSS (-)]** and collected after “***terminal death***” of HRC23 cells. Each kind of solution was carried out using three flasks for each solution (n=3). The error bar shows the standard deviation **±**SD in percentage obtained using the SDEVA function of MS-Excel. The figure is shown as sample concentration expressed as corresponding resource sample volume(ml)/ml in common logarithm (log_10_) on the horizontal axis, and the mean cell viability value of three flasks for each solution calculated from MTT assay in the percentage against non-treated control with an error bar as mean ±SD on the vertical axis. The SD values in the three graphs were at the worst point. **Non-fractioned: solid line (), fraction with MW<1 kDa: broken line (------)**.

### 2. Examination of nutritional environment on the cytocidal substance(s) production

At first, the effect of serum on **“*terminal death*”** was examined. At the “terminal phase,” the culture medium was replaced with a serum-free medium (this solution replacement with other types of solutions during the “terminal phase” is called the ‘final solution replacement method’ in this study) and incubated (**Figure 1. B**.) When serum-free EMEM was used as a final solution, the result showed **“*terminal death*”** and cytocidality with concentration dependency (**Figure 3. EMEM**.) So, separation of the resource sample depending on molecular size was examined. The collected serum-free EMEM was separated into 2 fractions with an MW of 1kDa in between by ultrafiltration. The fraction of MW<1kDa **(Figure. 3: EMEM<1kDa)** was similar to the case of serum-containing culture medium **(Figure 3: EMEM + FBS.)** As the fraction with MW>1kDa did not exsert cytocidality, the size of cytocidal exserting substance was suggested to MW <1kDa.

Next, to examine whether this cytocidal substance(s) production depends on the nutrients and energy resources contained in the medium, the ‘Final Solution Replacement Method’ was performed with glucose-free HBSS [shown as HBSS (-).] We call this method a **‘Nutrition-Free Method’**. Surprisingly, in this ‘**Nutrition-Free Method**’, *“****terminal death****”* was also recognized and detected cytocidality with concentration dependency in the fraction with MW<1kDa of the supernatant obtained by ultrafiltration as in the case of serum-free medium (**Figures 2. and 3**.) The results of this **‘Nutrition-Free Method’** showed that when cells enter from the “terminal phase” to the “***terminal death***,” they produce a cytocidal substance(s) even in an environment without nutrient and energy sources. Furthermore, since the cytocidal sample obtained by this ‘**Nutrition-Free Method**’ was considered to be produced by cells only from intracellular component(s), it was also important to make cytocidal substance(s) purification far easier.

### 3. Cytocidal effects CSS produced by HRC23 against other mammal cancer cell lines

The cytocidal effect of the first gel filtration processed CSS sample produced by HRC23 against four other histologically different pattern human cancer cell lines and mouse <u>L</u>ewis <u>L</u>ung <u>C</u>arcinoma (LLC) ^4^ was examined (**Figure 4**.) HRC produced CSS exserted cytocidality against itself and tested the other five cell lines regardless of differences in histological patterns, as epithelial or non-epithelial origins and species. The pattern of dose-response and morphological changes during cell death caused by CSS was common to all the cell lines tested. The fact that a substance produced by one type of cell exerted similar cytocidal effects in the dose dependency suggests that they have the same system that reacts to CSS. Furthermore, as the “***terminal death***” was confirmed in all cells examined, the production of CSS, or a similar substance, by these cells at the ‘terminal phase’ is possible. It was noteworthy that HRC23 (a human cancer cell) produced CSS killed even different species of cancer cell line (mouse LLC) as others. So, the cytocidality of CSS produced by a cancer cell line against other types of cancer cell lines (**Table 1**.) was investigated. (**Figures 4**.,**5. and 6**.) 3. Cytocidal effect of the whole and its fraction with MW<1kDa of HBSS(-) obtained after cancer cells death using ‘**Nutrition-Free Method**’ against themselves, respectively.

**Table 1.**

| **Cell name**<br>**(code number)** | **Species**<br>**Organ** | **Histological type**<br>**of original tumor** | **Culture**<br>**Medium** | **Trypsin**<br>**and EDTA**<br>**concentration.** |
| --- | --- | --- | --- | --- |
| **HRC23 (original,**<br>**Same to ku-2)<sup>3, 4, 5)</sup>** | **Human**<br>**Kidney** | **Adenocarcinoma** | **EMEM**<br>**with 10%**<br>**FBS** | **0.1% T**<br>**0.01% E** |
| **MKN74**<br>**(JCRB0255)** | **Human**<br>**Stomach** | **Adenocarcinoma**<br>**(tubular)** | **RPMI1640**<br>**with**<br>**10% FBS** | **0.25% T**<br>**0.02% E** |
| **LK-2 (JCRB0829)** | **Human**<br>**Lung** | **Squamous cell**<br>**carcinoma** | **RPMI1640**<br>**with 10%**<br>**FBS** | **0.125% T**<br>**0.01% E** |
| **VMRC-JCP**<br>**(JCRB0103)** | **Human**<br>**Lung** | **Squamous cell**<br>**carcinoma** | **RPMI1640**<br>**with 10%**<br>**FBS** | **0.125% T**<br>**0.01% E** |
| **SKN**<br>**(JCRB0173)<sup>1)</sup>** | **Human**<br>**Uterus** | **Leiomyosarcoma** | **EMEM**<br>**with 10%**<br>**FBS** | **0.125% T**<br>**0.01% E** |
| **LLC (RBC0558)<sup>2)</sup>** | **Mouse**<br>**Lung** | **Medullary**<br>**carcinoma** | **EMEM**<br>**with 10%**<br>**FBS** | **0.04% T**<br>**0.003% E** |
| **BHK, BHK-21(C-13)**<br>**(JCRB9020)** | **hamster,**<br>**Syrian,**<br>**kidney,**<br>**subclone of**<br>**BHK-21** | **fibroblast-like** | **EMEM**<br>**with 10%**<br>**calf serum.** | **0.25% T**<br>**0.02% E** |
| **Vero**<br>**(JCRB0111)** | **Monkey,**<br>**African**<br>**green,**<br>**kidney** | **epithelial-like** | **EMEM**<br>**with 5%**<br>**FBS** | **0.25 % T**<br>**0.02 % E** |

**The code number in the bracket of cells is peculiar to distributed institutions.** **JCRB: Japanese Collection of Research Bioresources Cell Bank (JCRB Cell Bank; National Institutes of Biomedical Innovation, Health and Nutrition, Osaka, Japan).** **RBC: RIKEN BRC, “RIKEN BioResource Research Center. “(Tsukuba, Ibaraki, Japan)**

**Figure 4.**
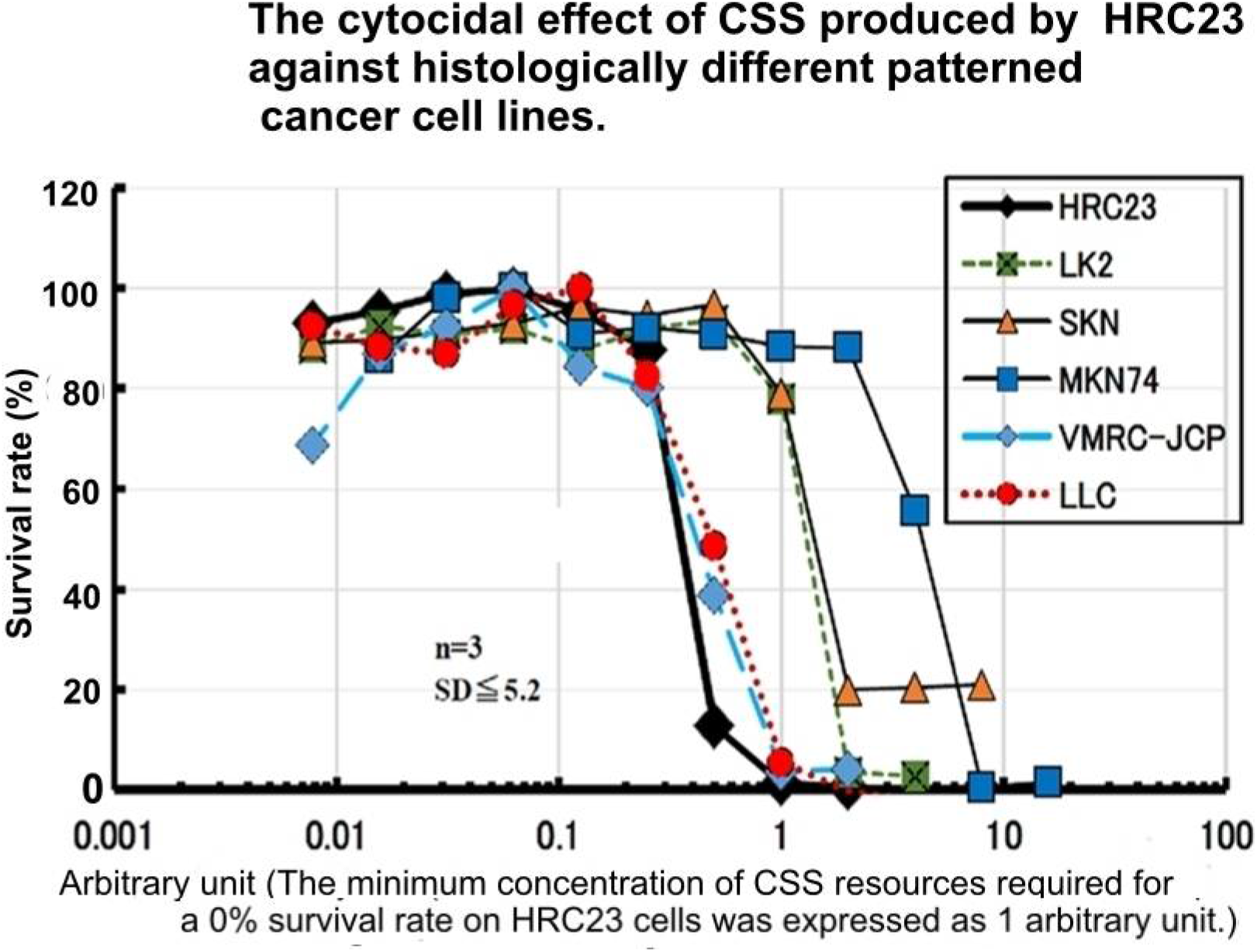
The cytocidal effect of CSS produced by HRC23 against histologically different patterned cancer cell lines (Table 1.) HRC23 cells were employed as a reference to correct the difference of each cell assay plate. The CSS sample concentration [in a two-fold series dilution concentration was resource sample volume (ml/ml)] where HRC23 (reference) viability becomes 0% is defined as the arbitrary unit as Arbitrary unit 1. Viability was measured by the MTT assay method. In the semi-log graph, the cell survival rate is shown with mean ± SD value in percentage of the non-treated control. The marker and line colors for the cells are: HRC23 (reference): 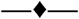, LK2: 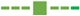, SKN: 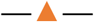 MKN74: 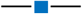, VMRC-JCP: 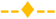, and LLC: 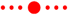

**Figure 5.**
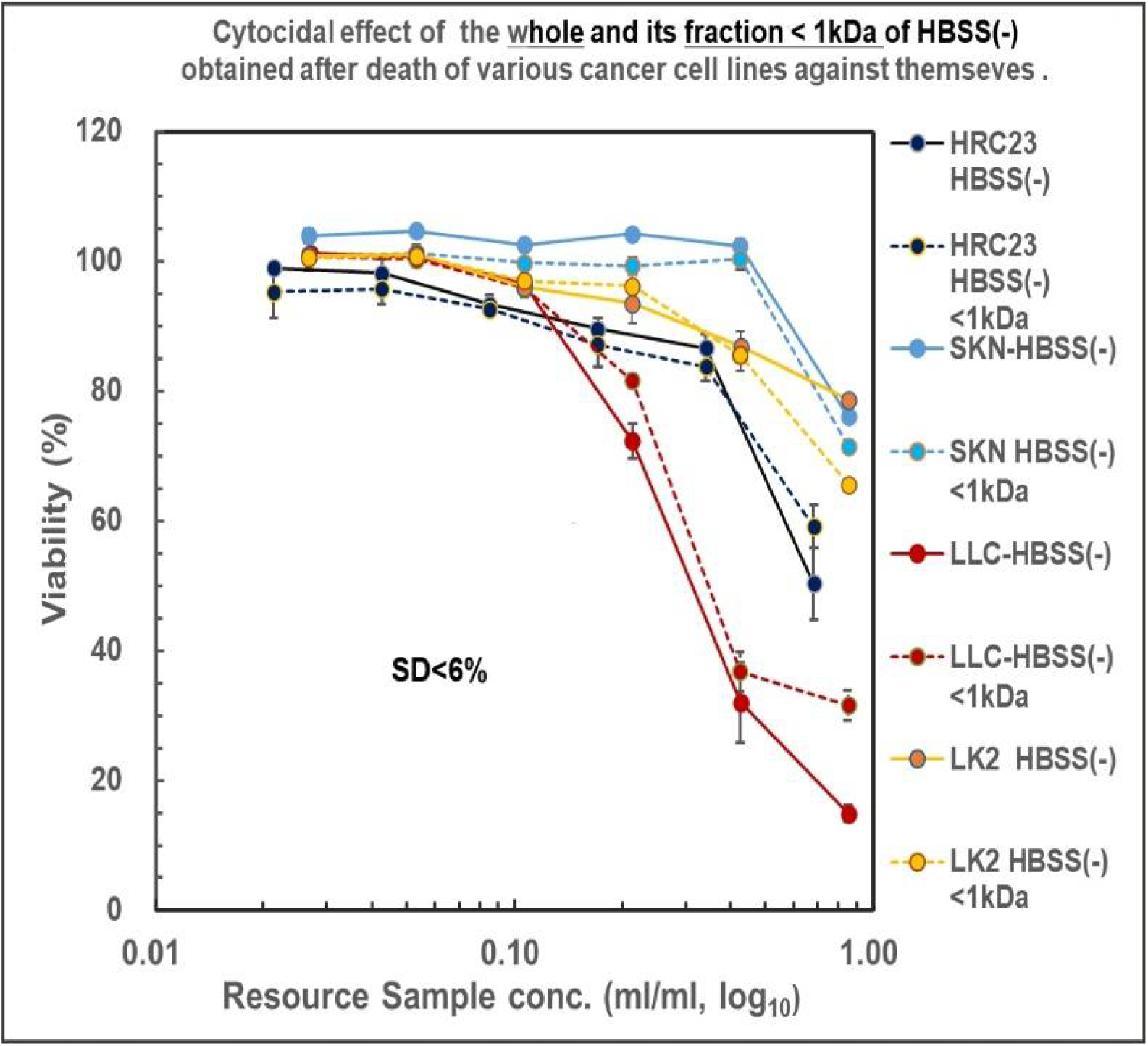
Cytocidal effect of the HBSS(-) and its fraction (MW<1kDa) obtained by HRC23, LK2, SKN and LLC using ‘**Nutrition-Free Method**’ against the cell lines themselves, respectively. The samples were regenerated, and the viability were measured according to the ‘resource sample assay procedure’^*1^ by the MTT assay method. In the semi-log_10_ graph, the cell survival rate was expressed with mean ± SD in percentage for the non-treated control. Marker and line colors for the cells are: **(—)filled line: HBSS (-), (-----)dotted line: HBSS(-)<1kDa**. 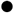: **HRC23**, 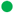: **LK2**, 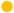: **SKN**, 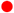: **LLC. *1: Shown in Detailed Method 2-3**

**Figure 6.**
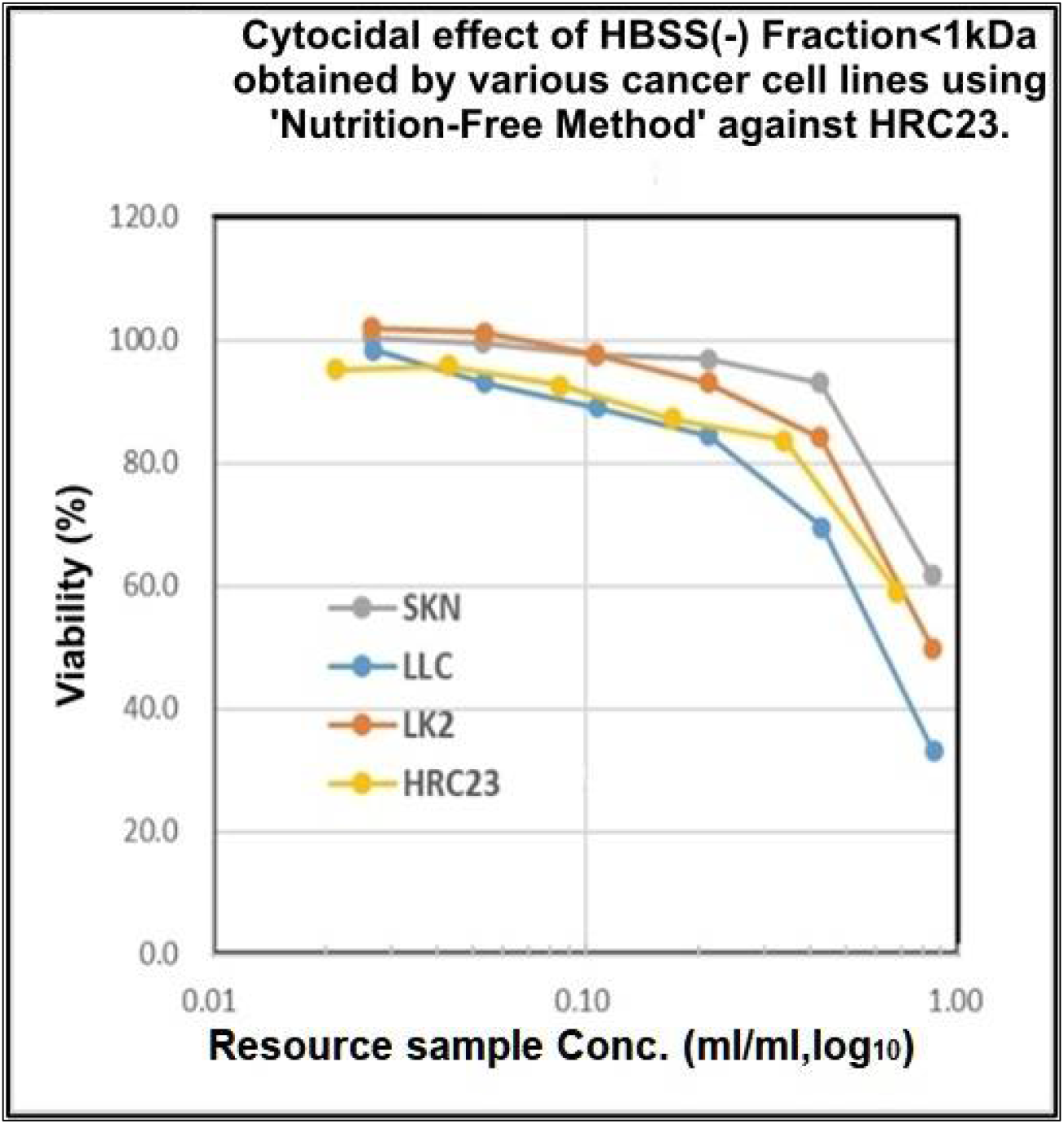
Cytocidal effect of the fraction with MW<1kDa obtained as of **Figure 5**. Against **HRC23**. The samples were regenerated, and the viability were measured according to the ‘resource sample assay procedure^\***2**^by the MTT assay method. In the semi-log graph, the cell survival rate (vertical axis) was expressed with mean ± SEM in the percentage of the non-treated control. Marker and line colors for the cells are 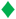: **HRC23**, 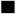: **LK2**, 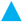: **SKN**, 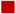: **LLC**. \***2** Shown in Detailed Method 2-3

On HRC23, SKN, LLC, and LK-2 cells, the cytocidality of CSS produced by each cell line against themselves was investigated. Both fractions (whole and <1kDa) of the four all-cell lines exserted cytocidality against themselves with dose dependency. Hence cytocidal substance was suggested to be contained in the fraction with MW<1kDa for all cancer cell lines tested.

### 4. Cytocidal effect of HBSS (-) fraction with MW<1kDa obtained by the other cancer cell lines against themselves and HRC23

As cytocidal substance(s) existed in the fraction with MW < 1 kDa, the cytocidality of CSS produced by each cell line against HRC23 was investigated. The fraction of all cell lines exserted cytocidality against HRC23 with dose-dependency (**Figure 6**.) These results (**Figure 5. and Figure 6**.) showed the CSS production by cancer cells exert cytocidality against themselves and other cancer cells and vice versa. Now CSS production by cancer cells has been confirmed.

### 5. The cytocidal effect was not recognized in the solution obtained by nontumor cell lines using the ‘Nutrition-Free Method’

As the nontumor cell line, BHK and Vero cell lines were employed. After the cell death, which is caused by starvation but not *“****terminal death***,*”* the whole solution, and the fraction with MW <1kDa were prepared and regenerated for the cytocidality assay. Both cell lines showed no cytocidality against themselves (**Figure 7**.) and HRC23 (**Figure 8**.) These results led to CSS production as a specific event for cancer cells but not for nontumor cells. CSS exerts cytocidality against its cells itself and other cancer cells despite histologically different patterns and species.

**Figure 7.**
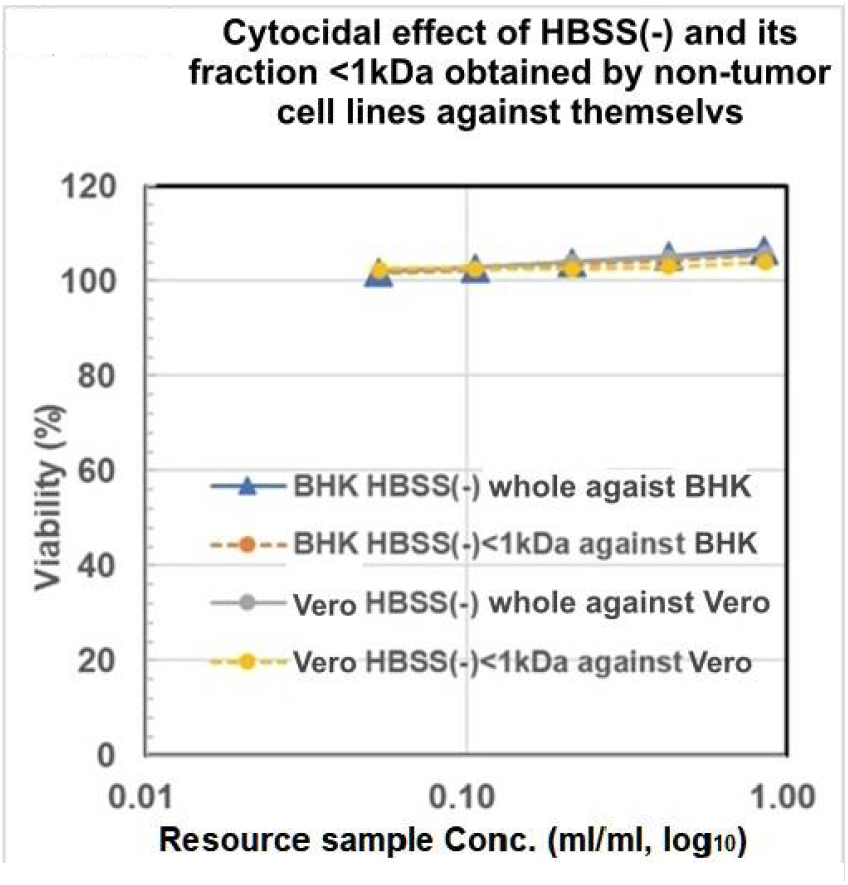
Cytocidal assay of the resource sample made by Nontumor cell lines against themselves. Cytocidal effect of the hole and fraction <1kDa of HBSS (-) obtained after the cell death (“**Nutrition-Free Method**”) against themselves, respectively. Viability was measured by the MTT assay method. Sample concentration is shown as resource sample volume (ml/ml) in log_10_ on the horizontal axis, cell survival rate was expressed with mean ± SD (<3%, n=3) in the percentage of the non-treated control on the vertical axis. Marker and line colors for the cells are BHK HBSS (-) whole: 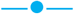, BHK HBSS(-)<1kDa: 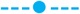, Vero HBSS(-) whole: 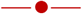, and Vero HBSS(-)<1kDa: 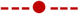.

**Figure 8.**
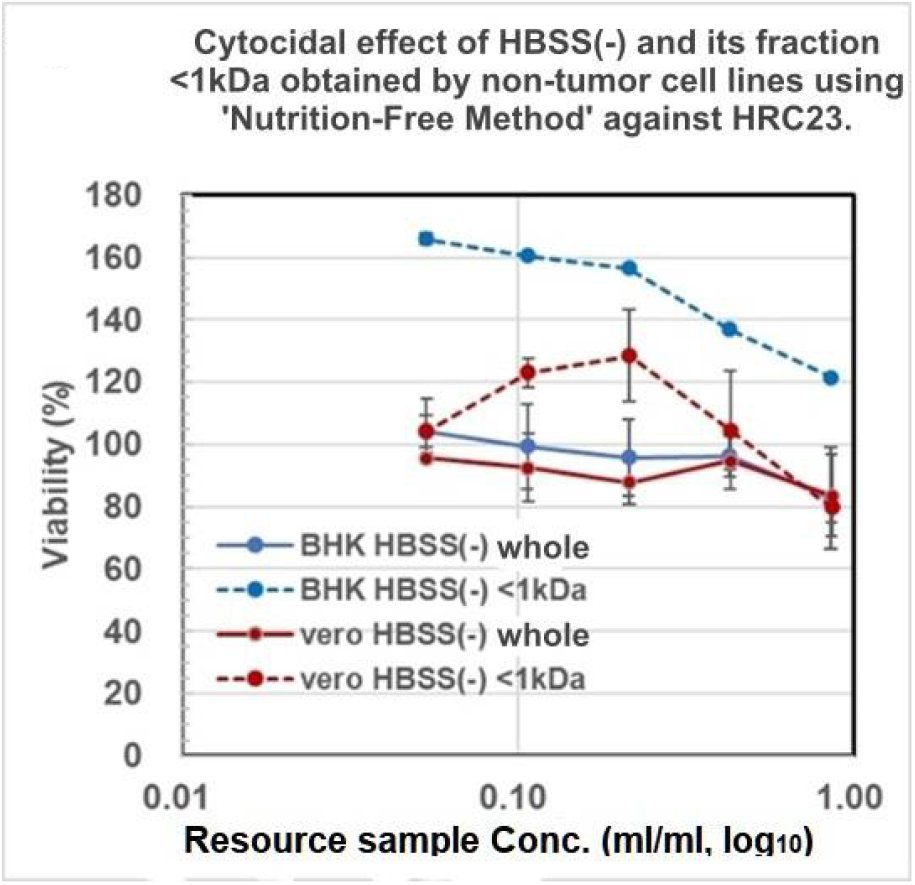
Cytocidal assay of the resource sample made by Nontumor cell lines against HRC23. Cytocidal effect of the resource sample by BHK and Vero obtained using ‘N**utrition-Free Method**.’ The regenerated HBSS (-) and its fraction with MW<1kDa obtained as of **Figure 7** against HRC23. Viability was measured by the MTT assay method. Sample concentration is shown as resource sample volume (ml/ml) in log_10_ on horizontal axis, cell survival rate was expressed with mean value in percentage of the non-treated control with error bar of ±SD (<7%, n=3.) Marker and line colors for the cells are BHK HBSS(-) whole: 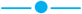, BHK HBSS(-)<1kDa: 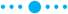, Vero HBSS(-) whole: 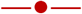, and Vero HBSS(-)<1kDa: 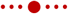.

### 6. Effect of CSS on LLC-transplanted mice

Thus, CSS killed cancer cells regardless of histologically different patterns and even distinct species *in vitro*. To investigate the anticancer effect of CSS *in vivo*, the first gel filtration processed HRC23-produced CSS was administered into LLC-transplanted mice. Two control mice died simultaneously on the 25th day after transplantation; however, CSS-administered mice survived longer than the control. CSS administered one mouse died on the 35th day and another mouse died on the 48th day after transplantation (**Figure 9**.) Hence CSS prolonged CSS-administered mice life spans clearly, compared to the control group. In addition, adverse events were not observed during the 6 days of administration with the CSS.

**Figure 9.**
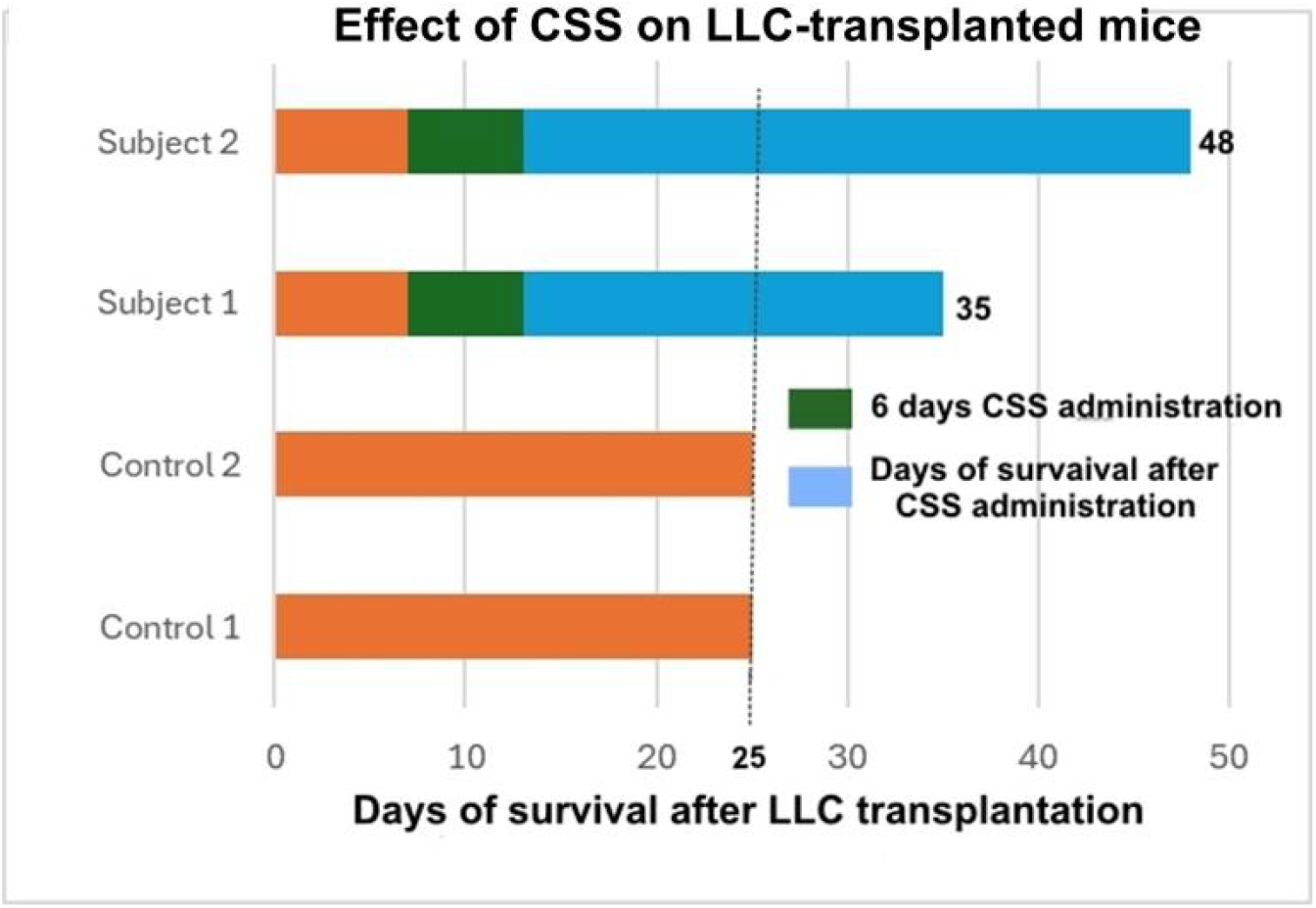
Effect of CSS on LLC-transplanted mice. LLC cells (3×10^6^ cells/mouse) were transplanted intraperitoneally into four mice (at 6 weeks of age). One week after LLC transplantation, gel filtration processed “CSS” sample, equivalent to 1000ml resource sample of HRC23 prepared by **‘Nutrition-Free Method’** was injected intraperitoneally into the two mice each, once a day, for six days. Another 2 mice were used as non-CSS treated control. In the graph, green colored block shows CSS administration period. Blue colored block shows survived period after CSS administration.

### 7. Purification

The extracted fraction with ethanol from dried resource sample solution obtained by the ‘Nutrition-Free Method’ of HRC23 cells was further purified after EtOH extraction by gel filtration column chromatography (**Figures 11. A and C**) and strong-cation exchange column chromatography (**Figure B**.) At first, gel filtration column chromatography was performed (**Figure 11. A**.) The cytocidal effect was detected only for the fractions of group **<u>A</u>** (filled fractions between 105 min to 141 min in (**Figure 11. A**,) just after the void volume (column gel MW exclusion size 1 kDa, designated as V_0_), with almost 100% activity recovery. The first gel filtration processed CSS sample exhibited a steep change in the cytocidal assay by an MTT method; the concentration range where cell viability changes from 100% to 0% by the two-fold concentration of CSS sample in (**Figure 10. C**) which was visibly obvious on MTT assay plate image (**Figure 10. B**) than that of the resource samples (**Figure 3**., **5. And 6**.) In addition, the cells were destroyed into small fragments (**Figure 10. A:right image**). Next, the cytocidal effect detected fractions obtained by first gel filtration were applied to a strong cation exchange column and eluted by the linear gradient method on Na_2_SO_4_ concentration (**Figure 11. B**.) The cytocidal effect exhibited fraction was detected at the valley region (in the fraction eluted between 165 mM to 170 mM Na_2_SO_4_, filled fractions in the chart) on the chromatogram, yielding approximately 10% activity of the extracted sample with ethanol. As the cytocidality exhibited fraction was in the valley of the elution profile, second gel filtration column chromatography was performed for the fractions (**Figure 11. C**,) where the elution profile was monitored by absorption at 205 nm instead of at 230 nm as specific absorbance band was not found. One large peak with a cytocidal effect in the first front of the chromatogram and a middle peak without cytocidality were detected. To verify the purity of the cytocidal effect detected fractions, those were divided into three fractions (designated as Fr.57, Fr.58, and Fr.59 in the chromatogram) and supplied for the TOF-MS analysis (**Figure 12**.)

**Figure 10.**
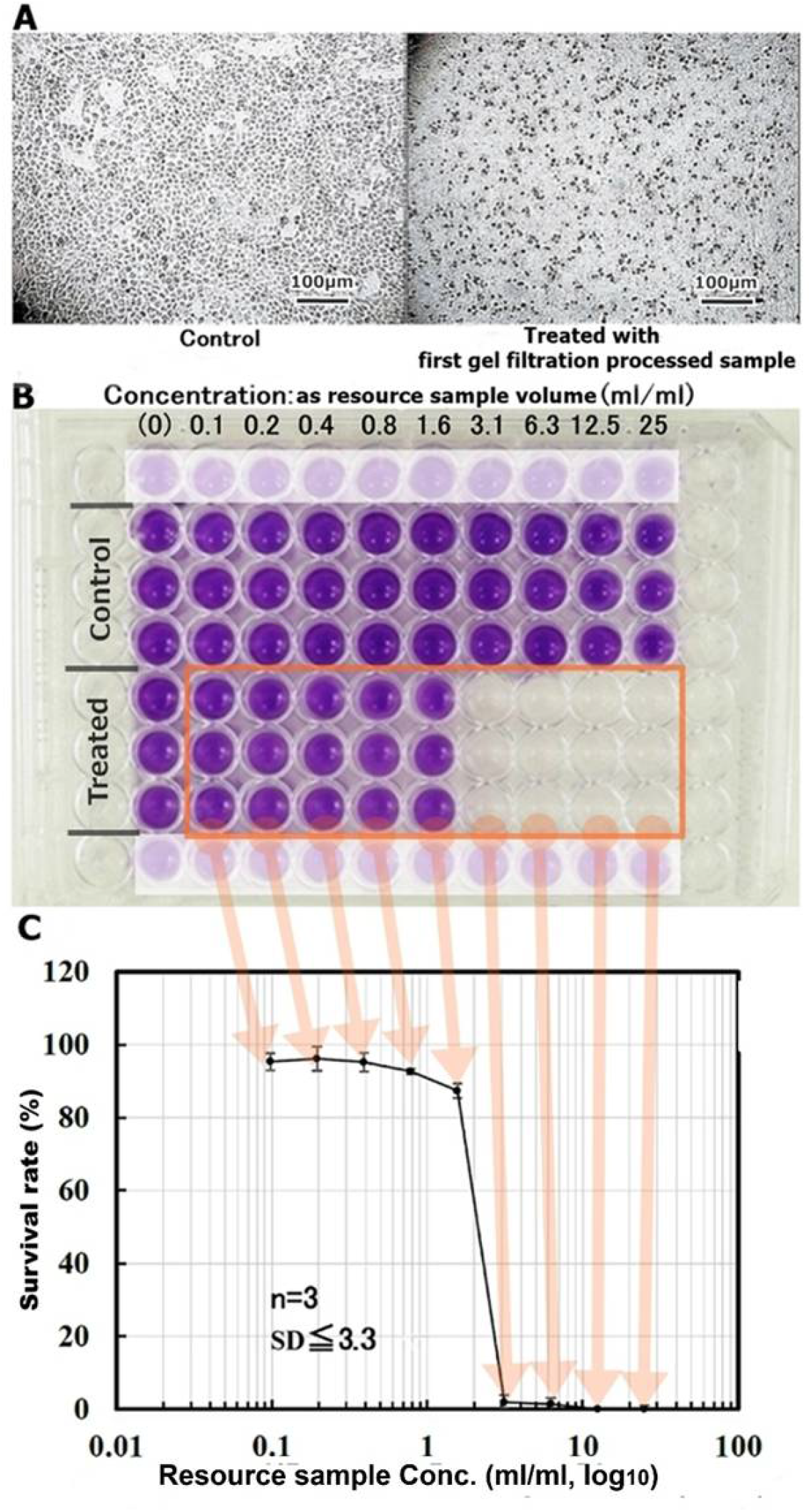
Cytocidal assay of first gel filtration processed CSS sample. **A: Microscopic image of the cytocidal effect of gel filtration column chromatography-processed CSS sample against HRC23 cells**. At CSS resource sample concentration was 3.1ml/ml, viability was 0 % (right image), and untreated control (left image). The inserted bar indicates 100µm in both culture images. **B: Image of MTT assay using 96 well microplate on the cytocidal effect of the first gel filtration-processed CSS sample against HRC23 cells**. A series of 2-fold dilutions CSS sample were replaced for the regenerated medium of 1 day precultured HRC23. After two days of culture with the sample, cell viability was measured by an MTT assay method. **C: Cytocidal effect of the first gel filtration chromatography-processed CSS sample on HRC23 cells using an MTT assay**. This semilogarithmic graph was deduced from the photometry of the MTT assay microplate in (**Figure 6. B.)** CSS sample concentration was expressed as corresponding resource sample volume (ml) /ml by common logarithm (base 10) on the horizontal axis and mean cell viability value calculated from MTT assay in the percentage against non-treated control with an error bar as mean ± SD on the vertical axis.

**Figure 11.**
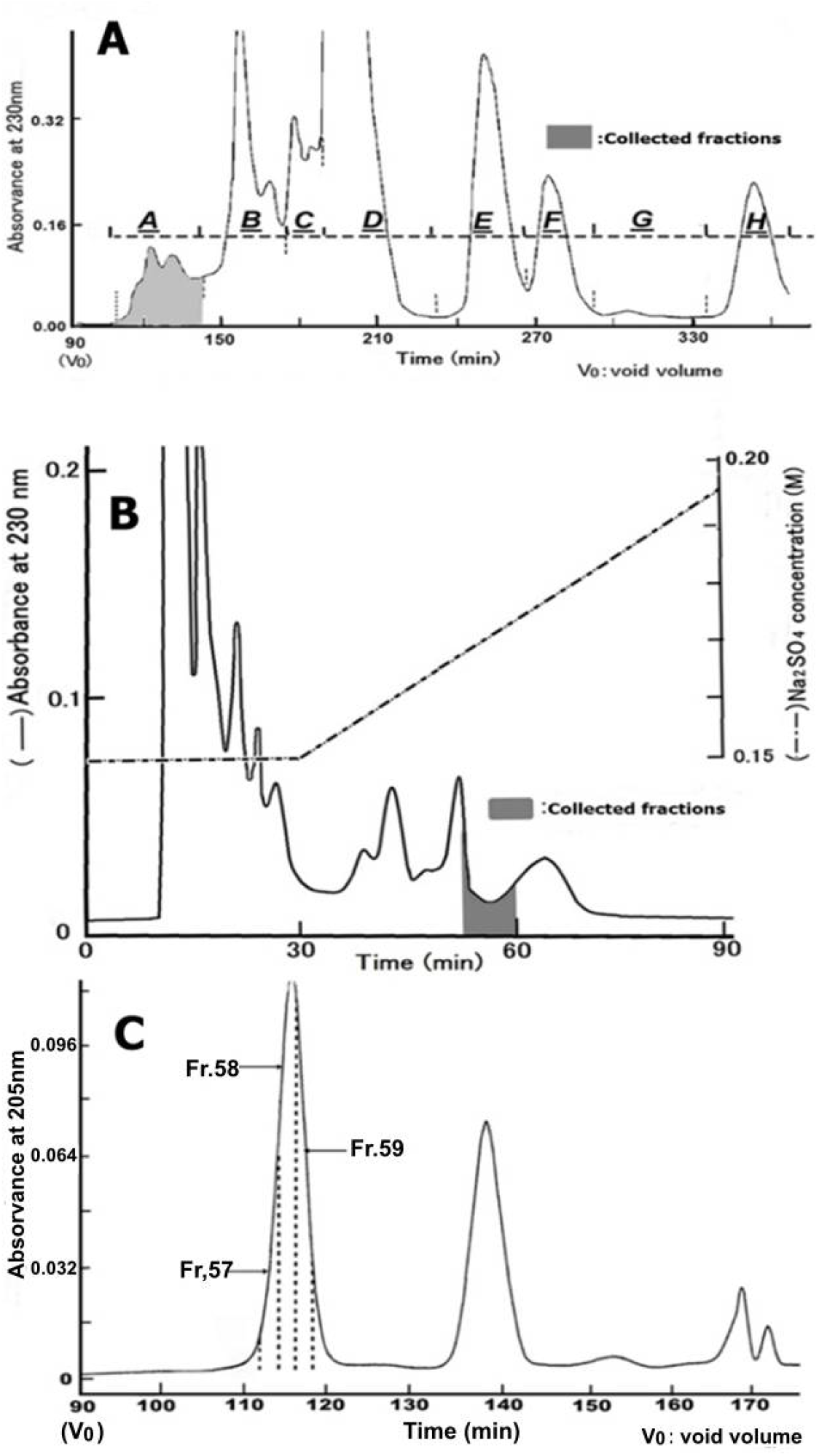
CSS purification by column chromatography. **A: Gel filtration column chromatogram**. The dried resource sample, which was prepared from 2 liters of resource sample solution obtained after the death of HRC23, cultured with 600 HBSS (-) as the final solution 601 replacement, was applied. Eluted 602 fractions were divided into 8 603 groups designated as **<u>A</u> t**o **<u>H</u>**. Only group **<u>A</u> (**filled fractions) exhibited a cytocidal effect. **B: Ion (cation) exchange column chromatogram**. The cytocidality exhibited fraction group (A) obtained by first gel filtration was used. An aliquot of 611 the fraction was used for cytocidal activity assays using the MTT method. Cytocidality exhibited fractions (filled fractions on the chromatogram) were pooled and dried after removal of Na_2_SO_4_ by adding the same volume of MeOH or EtOH, 619 then supplied for the second gel filtration column chromatography. **C: Second gel filtration column chromatogram**. Cytocidal effect exhibited fractions obtained by the ion exchange column chromatography was applied and eluted as in **(Figure 4. A.)** The cytocidal effect exhibited fraction was present only in the peak at first in front of the chromatogram. To know the purity of the fractions, they were divided into three parts denoted as Fr.57, Fr.58, and Fr.59. After the removal of Na_2_SO_4_, the three fractions were dried and used for TOF-MS analysis

**Figure 12.**
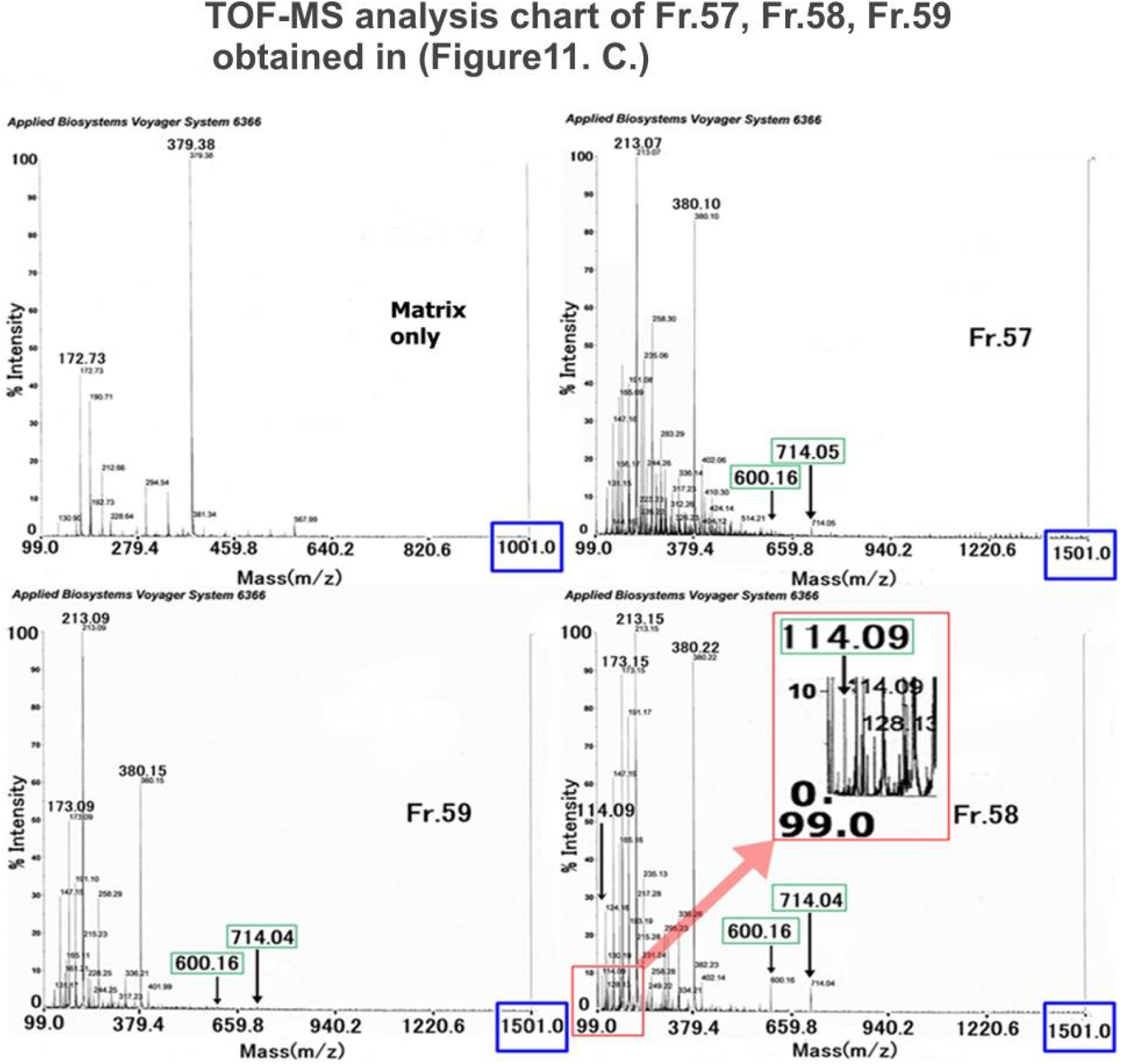
TOF-MS analysis. TOF-MS analysis spectra of the three fractions (Fr.57, Fr.58, and Fr.59) obtained by the second gel filtration column chromatography **(Figure 11. C.)** In the spectrum of Fr.58, the inserted spectrum is the enlarged figure around the m/z 114.09 signal. M/z 714.04 and m/z 600.16 signals were detected for the tree fractions, but larger than m/z 714.04(5) signal was not observed.

### TOF-MS analysis

Obtained Fr.67, Fr.58 and Fr.59 by second gel filtration column chromatography were analyzed by TOF-MS. The m/z value measured was between m/z 100 and m/z 1500 (blank: matrix only was between m/z 100 and m/z 1000). In the spectra, between m/z 300 and m/z 1000, two signals at m/z 714.34 and m/z 600.47 were observed for all three groups. In addition, a signal at m/z 114.09 was observed clearly in the Fr.58 spectrum (inserted enlarged spectrum.) However, no signal was greater than m/z 1000.

## Discussion

Cancer cell lines can be cultured infinitely by passage, but suddenly they die without passage. Popularly this cell death has been considered to be caused by nutrition deficiencies, imbalances in the medium and etc. If the timing of passage was delayed, this cell death **“*terminal death****”* can be generally recognized, despite the medium being replaced at last evening with a fresh one. We thought that this cell death does not necessarily occur in the nutritional environment alone. Therefore, under the hypothesis that the medium after cell death (“***terminal death***”) contains certain cytocidal substance(s), this study was performed to make clear that “***terminal death***” was caused by some kinds of cytocidal substance(s) which may be produced in the cells in “terminal phase.” This study revealed that the water-soluble, cationic, and single small-size molecule with an MW<1kDa exserting cytocidality in the medium after **“*terminal death*”. “*Terminal death*”** was observed regardless of the nutritional environment and even in a nutrition-free physiological saline solution: HBSS (-). CSS was suggested to be produced only from “*<u>intra</u>cellular*” materials between the “terminal phase” and “***terminal death***” as a function automatically programmed into the cell itself. Judging from the result that nutritional environment as an “*<u>extra</u>cellular*” factor was independent of CSS production, and a certain “*<u>inter</u>cellular*” factor at the “terminal phase” is inevitably thought to be the initial cause to induce “*<u>intra</u>cellular*” CSS production and kills own cell, so cells produce a self-killing substance (CSS) initiated by a certain change of “*<u>inter</u>cellular*” condition. “***Terminal death***” can be recognized as a type of physiologically programmed cell death. This physiological substance is produced by its physiological process and kills its cells. CSS can be the direct and conclusive substance that results in ***“terminal death*.”** The **‘Nutrition-Free Method’** promoted CSS purification extremely. Also, the method contributed to further study on other cell lines that will be seen soon. The chemical formula and conformation analysis of the CSS is in progress. Some conditional changes between cells concerning CSS production are still unknown. While currently used anticancer drugs are difficult to dissolve in aqueous solutions, it will be noteworthy that CSS is highly water-soluble with a molecular weight of less than 1kDa substance. This study showed that CSS production is a cause of “***terminal death***” and specific events of cancer cells, and cells have a common system to react to CSS. CSS can be called a suicidal factor of a cancer cell even in a monolayer system, so we hope to consider cancer therapy.

### 1. CSS production does not depend on extracellular nutritional conditions

It was necessary to investigate how the nutritional environment acts as an extracellular factor for the appearance of “***terminal death***,” and whether biologically active substances, produced in culture, caused the growth arrest of cells during overpopulation in the terminal phase as an intracellular factor. **At first**, to reproduce *“****terminal death***,*”* which is always seen in our laboratory after culture was renounced passage, a final solution exchange was performed with FBS added medium for HRC23 culture. Confirming that the culture was in the *“****terminal death****”* phase by microscopic observation, the medium was collected. This supernatant of the collected medium was regenerated and applied to HRC23 cultures. The result showed that the collected medium had a cytocidal effect and exhibited a dose dependency against HRC23 cells. This phenomenon suggests that the collected medium contained a cytocidal substance against the cells from which it was derived. Therefore, we named this cytocidal substance: <u>c</u>ancer <u>c</u>ell <u>s</u>uicide <u>s</u>ubstance (CSS).

**Secondly**, to verify whether serum is necessary for CSS production, a CSS production experiment was performed with serum-free EMEM at the ‘final solution exchange.’ The results were similar to those obtained with an FBS-containing case. To confirm that CSS was contained in the cell-dead solution, and to examine the molecular size of CSS, ultrafiltration was done with a MW cut-off membrane filtration at 1kDa. The fraction with MW<1kDa, the same cytocidal effect, and dose dependency against HRC23 cells were confirmed. The fraction with MW>1kDa did not exhibit cytocidal effect. Thus, the MW of CSS is suggested to be a MW<1kDa substance and serum is unnecessary for CSS production.

**Thirdly**, to examine the necessity of nutrition for CSS production, a final ‘solution exchange’ was performed with glucose-free HBSS(-). When HRC23 was incubated with this solution, the culture also displayed ***“terminal death*.*”*** The sample obtained in this experiment showed the same effect as the former experiments, specifically, the dose-dependency of the cytocidal effect was exhibited by the fraction with MW<1kDa. The latter two results suggested CSS to be a low molecular weight substance with MW<1kDa. In addition, CSS is produced only from intracellular materials, and CSS production is independent of the nutritional environment.

### 2. The relationship between CSS production and apoptosis

Based on the result that cells produce a self-killing substance, independent of external nutrition, *“****terminal death****”* can be recognized as a type of physiological cell death. In addition, CSS production is activated by intracellular changes that occur during specific conditions known as the “terminal phase,” and this substance kills its cells. This phenomenon seemed remarkably similar to the concept of apoptosis.^10, 11^ ^12, 13.^

### 3. The effect of CSS against other cancer cell lines, and the possibility of applying CSS as an anti-cancer therapy

HRC23-produced CSS kills not only against HRC23 cells, from which it was produced, but also against the other types of cancer cells regardless of differences in histological patterns, such as epithelial or non-epithelial origins, or even species. The pattern of dose-response and morphological changes during cell death caused by CSS was common to all the cells tested. The fact that a substance produced by one type of cell exerted similar cytocidal effects against all five histologically different pattern cells examined, suggests that they have the same system that reacts to CSS. This fact indicates that the mechanism of the cytocidal effect of CSS is unique and different from that of known substances. On benign tumor cells, we could not obtain the cell line.

### 4. Effect of CSS on LLC-transplanted mice

CSS kills many cell lines derived from other histologically distinct types of malignant neoplasms *in vitro*. However, the question remains whether CSS is effective *in vivo*. We preliminarily tested the effect of CSS on LLC-transplanted mice. LLC is a cell line of mouse cancer origin. LLC is reported to be resistant to many kinds of antitumor drugs. ^3^ The result showed a clear difference in life span by suppression for LLC growth without any adverse events during the CSS administration period: between CSS-treated and non-treated control. Thus, CSS has potential as an anticancer drug.

## Conclusion

Regarding spontaneous cell death of cancer cell lines, our study found that cancer cell lines produce a cytocidal substance CSS. CSS was found in the medium after cells died, and CSS was made even in a nutrition-free physiological solution. Therefore, CSS was produced only from the intracellular components themselves by the cell itself and killed its cell. CSS is made by other histologically different cancer cell lines regardless of species of mammals and vice versa. CSS was a small molecule with an MW<1kDa and highly water-solved character in contrast to currently used anticancer drugs. In addition, CSS was considered as the direct and conclusive substance that results in **“*terminal death*.” “*Terminal death*”** of cancer cell lines is commonly known as one of the forms of physiological cell death. CSS is produced “*<u>intra</u>cellularly*” caused by “*<u>inter</u>cellular*” condition change in the “terminal phase.” CSS production should be a physiologically programmed phenomenon. This phenomenon can be understood as a cell function of multicellular systems. This phenomenon was aroused by a change in the ‘*<u>inter</u>cellular*’ community which can be understood as a cell function of multicellular systems.^16^ *In vivo* examination, it is noteworthy that CSS suppressed metastasis and prolonged the life of LLC-transplanted mice without any adverse events. Hence CSS has the possibility for a *‘new type’* of anti-cancer agent development. We hope this study contributes to further studies given new aspects of cancer therapy and would like to present this study as an example of an *“<u>inter</u>cellular”* community.

## Abbreviations

EMEM: (Eagle’s Minimum Essential Medium)
DMSO: (dimethyl sulfonic oxide)
HBSS(-): (Hank’s balanced saline solution without glucose)
FBS: (fetal bovine serum)
EtOH: (ethyl alcohol)
EDTA: (ethylene diamine tri acetic acid)
TOF-MS: (matrix-assisted laser desorption/ionization time-of-flight mass spectrometry)
MTT: [3-(4,5-dimethylthiazol-2-yl)-2,5-diphenyltetrazolium bromide]
TFA: (trifluoroacetic acid)
MW: (molecular weight.)

## Declarations

### Author Agreement

We authors have seen and approved the last version of the manuscript being submitted.

### Conflict of Interest and Competing Interes

There is nothing to declare. The author’s agreement.

### Declaration of Interest

This study and manuscript are managed only by the income of “Medical Corporation Ichikawa Clinic.” There was no support other than the clinic.

### Funding Source Declaration

This study is funded, managed, and supported only by “Medical Corporation Ichikawa Clinic.”

### Permission Note

This study and manuscript are original and performed by the authors of T. Tajima and Y. Kondo only. There is no need to get permission besides the authors.

## Author Contributions: CRediT taxonomy

**Conceptualization**, T. Tajima; **Methodology**, T. Tajima, Y. Kondo; **Validation**, Y. Kondo; **Formal Analysis**. Y. Kondo; **Investigation**, T. Tajima, Y. Kondo; **Data Curation**, T. Tajima, Y. Kondo; **Writing – Original Draft**, T. Tajima, Y. Kondo; **Writing – Review & Editing**. T.Tajima, Y. Kondo; **Visualization**, T.Tajima, Y. Kondo; **Supervision**, T. Tajima; **Project Administration**, T. Tajima, Y. Kondo; **Funding Acquisition**, T. Tajima; **Resources**, T. Tajima

## Intellectual Property

### Disclose any patents or copyrights an author may have that are relevant to the work in the manuscript

- US11318162 <u>Espacenet – Bibliographic data</u> https://ie.espacenet.com/publicationDetails/biblio?CC=KR&NR=102483786B1&KC=B1&FT=D&ND=4&date=20221230&DB=EPODOC&locale=en_IE
- US11963978 <u>Espacenet - Bibliographic data</u> https://ie.espacenet.com/publicationDetails/biblio?CC=US&NR=11963978B2&KC=B2&FT=D&ND=4&date=20240423&DB=EPODOC&locale=en_IE
- EP3693467 https://ie.espacenet.com/publicationDetails/biblio?CC=EP&NR=3693467B1&KC=B1&FT=D&ND=4&date=20220803&DB=EPODOC&locale=en_IE
- EP4086354 https://ie.espacenet.com/publicationDetails/biblio?CC=EP&NR=4086354B1&KC=B1&FT=D&ND=4&date=20240501&DB=EPODOC&locale=en_IE
- JP 6673558 https://www.j-platpat.inpit.go.jp/c1801/PU/JP-6673558/15/ja
- JP 6861305 https://www.j-platpat.inpit.go.jp/c1801/PU/JP-6861305/15/ja

## Detailed methods

### 1.. Cell culture

An established cell line derived from <u>h</u>uman <u>r</u>enal cell <u>c</u>arcinoma-derived cells (HRC23), which we established (the same as ku-2 cell ^5, 6, 7^) from a tumor derived from human kidney cancer transplanted into nude mice at the Science Laboratory of Tokai University Hospital (Kanagawa, Japan) in 1983, was used. They were grown in Eagle’s MEM (EMEM) (Nissui Pharmaceutical Co. Ltd., Tokyo, Japan) supplemented with 10% fetal bovine serum (FBS) (HyClone™, Hyclone Laboratories, Inc., UT USA, and Tissue Culture Biological Inc., CA, USA) without antibiotics in a CO_2_ incubator (Model MCO-19AIC; SANYO Electronic Co. Ltd., Tokyo, Japan) under the conditions of 37 °C, 5% CO_2_, and saturated relative humidity. Trypsin (Trypsin 250, DIFCO™; BD Bioscience/Becton, Dickinson and Company, NJ USA) at 0.1% and 0.01% EDTA in Dulbecco’s phosphate buffered saline without calcium and magnesium (PBS-) were used for cell dispersion. The other cell lines used in this study are listed in the table (**S1: Table1**), and they were cultured according to the indicated conditions in the same manner as HRC23.

### 2.. Preparation of <u>c</u>ancer cell <u>s</u>uicide <u>s</u>ubstance (CSS) samples to examine extracellular nutritional conditions on the cytocidal effect using three kinds of solutions for the final solution replacement in HRC23 culture

#### 2-1.. Sample preparation

HRC23 cells were cultured until the “terminal phase” in a flask (25 cm^2^, Corning Incorporated, NY, USA.) To examine the nutritional condition for cytocidal substance production, the three kinds of solutions: A) 10%FBS-added EMEM, B) FBS-free EMEM, and C) glucose-free Hank’s balanced salt solution [HBSS (-)] were employed. For each solution, three flasks of HRC23 cells cultured until fully confluent phase were prepared for solution A, and six (three for the molecular weight cut-off fractionation) flasks for solutions B and C, respectively. The flasks were rinsed four times with 10ml/flask of each kind of solution over 7 hours incubation at 37℃, then replaced with the same solution at 5ml/flask for each solution, respectively. This solution replacement procedure is called ‘final solution replacement.’ After the final solution replacement, cultures were incubated until the cells died **(“*terminal death*.”**) Then the cultured solution after the cells died was collected from each flask as the three kinds of CSS resource samples, respectively. These CSS resource samples were centrifuged at 1000×g [KOKUSAN model: H-108N, equipped with swing rotor (parts # MT-030,) KOKUSAN Co. Ltd., Saitama, Japan. (https://kokusan.co.jp/discontinued)] and the supernatants were filtrated with a 0.1 μm membrane filter (Millex®-VV, MilliporeSigma, the life science business of Merck KGaA, Darmstadt, Germany) and stored in a sterilized plastic bottle at 4 °C, respectively (resource samples.) These experimental procedures are summarized in (**Figure 1. B)**. In the case of using HBSS (-) as the final replacement solution to make CSS resource samples, this method is designated as a **‘Nutrition-Free Method’** ^**17**. **18**. **19**. **20**^ in this study. For other cell lines listed in the **Table 1**., RPMI-1640 (cat#: R8758-1L, Merck, Sigma-Aldrich, see “specification sheet” or Certificates of Analysis(COA) were used. (https://www.sigmaaldrich.com/JP/ja/product/sigma/r8758#product-documentation), were used.

#### 2-2. Fractionation of sample solution by a membrane ultrafiltration method

The three flasks for each of the two kinds of sample solutions: serum-free EMEM and HBSS (-) were fractionated using a molecular weight cut-off membrane with a nominal molecular weight cut-off size of 1 kDa (Stirred Cell Model 8050 equipped with Ultracel® Amicon® YM1, Ultracel® ultrafiltration membrane PLAC04310, MilliporeSigma, the life science business of Merck, KGaA, Darmstadt Germany) respectively. The filtrate was stored in a sterilized plastic tube aseptically by passing through a 0.1-μm membrane filter (Millex® VV, Merck) at 4 °C.

#### 2-3.. Cytocidal effect assay of the resource samples

To optimize the nutritional conflict of collected three kinds of samples for cytocidal effect assay, 1/50 volume of 50-fold concentration of amino acids for EMEM (Cat# 16223004, Kohjin Bio, Saitama, Japan) and 100-fold concentration Vitamins for EMEM (Kohjin Bio, Saitama, Japan) were added followed by pH adjustment with 7.5% NaHCO_2_, then 1/100 volume of 200mM glutamine and 10% volume of FBS were added to the samples (*regeneration*). For RPMI1640 based resource sample, RPMI 1640 Amino Acids Solution (50×) (cat#: R7131-100ML, Merck, Sigma-Aldrich (https://www.sigmaaldrich.com/JP/en/product/sigma/r7131?icid=sharepdp-clipboard-copy-productdetailpage), RPMI 1640 Vitamins Solution (100×) (cat#: R7256-100ML, Merck, Sigma-Aldrich (https://www.sigmaaldrich.com/JP/en/product/sigma/r7256?icid=sharepdp-clipboard-copy-productdetailpage,) as same to EMEM. The samples were diluted in a series of eight steps of two-fold serial dilutions with culture medium, starting with the highest concentration in triplicate, using one standard 96-well microplate (Corning Incorporated, NY, USA) for each sample. For each sample, the medium of 1-day precultured cells was replaced with 170 μl of the diluted sample followed by twice exchanges with the freshly diluted same sample as first on days 3 and 5. The cell viability was measured on day 6 by an MTT assay method ^8, 9, 10^. MTT (Dojindo Laboratories, Kumamoto, Japan) was dissolved in PBS-without calcium and magnesium at a 10-fold concentration (5 mg/ml) and aseptically filtered with a 0.1-µm membrane and kept in a cryotube at 4 °C. After the culture was washed with 200 µl of culture medium, 150 µl of culture medium containing 0.5 mM MTT was added to each well. After incubating the assay plate for 30 to 40 min at 37 °C, the MTT solution was removed by suction and washed with 200 µl of culture medium. Then 200-µl of dimethyl sulfoxide (DMSO) was added to each well. The absorbance was measured using a microplate reader (Model 550, Bio-Rad Laboratories, Inc., CA, USA) at 570 nm. Cell viability was calculated and expressed as: Viability (%) ={[A_570nm_ (sample)]-[A_570nm_ (blank)]}/{[A_570nm_ (control)]-[A_570nm_ (blanc)]} ×100. Correct ion by absorbance at other wavelengths was not employed.

#### 2-4.. Measurement of the cytocidal effect of HRC23 produced CSS against other types of mammal cell lines (Table 1 and Figure 4.)

To examine the cytocidal effect of the first gel filtration processed HRC23 produced CSS sample against other malignant tumor-derived cell lines, five four other typical human cell lines: MKN74, LK-2, VMRC-JCP, SKN ^2^, were obtained from the <u>J</u>apanese <u>C</u>ollection of <u>R</u>esearch <u>B</u>ioresources Cell Bank (<u>JCRB</u> Cell Bank; National Institutes of Biomedical Innovation, Health and Nutrition, Osaka, Japan). Mouse <u>L</u>ewis <u>l</u>ung <u>c</u>arcinoma cell line (<u>LLC</u>; RCB0558), which is known to have enhanced lung metastatic ability and high resistance to all types of anti-cancer drugs,^15^ was also used [obtained from the <u>R</u>IKEN <u>C</u>ELL <u>B</u>ANK (<u>RBC</u>; Cell Engineering Division -Cell Bank-/ RIKEN BioResource Center, Tsukuba, Japan)]. Cell names, code numbers at the facilities, the organ of origin, histological types of the original tumor, culture medium, and trypsin-EDTA concentrations for cell dispersion are listed in **Table 1**. The dried HRC23 CSS sample was dissolved in a culture medium suited for the cells. After 1 day of preculture, the dilutions in a series of eight steps of two-fold serial dilutions with culture medium, starting from equal to 200ml of resource sample/ml. The culture incubation condition was the same as of HRC23.

#### 2-5.. Cytocidality investigation on nontumor cell lines

As a nontumor mammal cell line, two cell lines were employed. **BHK**: BHK-21(C-13) (JCRB9020) hamster, Syrian, kidney, subclone of BHK-21, and **Vero** (JCRB0111) Monkey, African green, kidney. The ‘**Nutrition-Free Method**’ was used for this investigation. The cytocidal assay was denoted in former sections **2-3 and 2-6: Cytocidal effect assay**.

#### 2-6.. Cytocidal effect assay of the EtOH-extracts of dried resource sample and the fractions of column chromatography processed CSS samples

HRC23 cells were inoculated onto a standard 96-well microplate separately (day 0) as HRC23 to be approximately 80% confluent in three days. After one day of preculture (day 1,) the culture medium was replaced with 170 μl of the two-fold dilution series samples for one well. The dilutions were prepared just before use on day 1 as follows: The dried CSS sample was dissolved in the culture medium and diluted by eight steps of a two-fold dilution series with culture medium, beginning from equal to 200 ml of resource sample /ml in triplicate in a standard 96-well microplate. After a further 2 days of culture with the sample, the survival rate of the cells was measured by the MTT assay method (day 3) as shown in above section **2-3: cytocidality assay**.

#### 2-7.. Statistics

All cytocidality assays were performed in triplicate independently. The mean value and standard deviation (SD) of viability were calculated using Microsoft Excel functions of “AVERAGE” and “STDEVA*^1^” respectively. The results were shown as mean value (marker) ±SD (error bar) on the figures. As the cytocidal effect measurement of CSS samples against the cells by an MTT assay was very reproducible (at worst≦6.5%,) MTT assay size for each independent measurement was done in triplicate. *^1^ STDEVA uses the following formula: 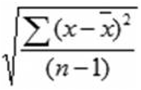 where x is the sample mean AVERAGE (value1, value2…) and n is the sample size.

### 3.. CSS purification

#### 3-1.. Sample preparation by a large-scale culture

The flask with a 175cm^2^ culture area (Greiner Bio-One International GmbH, Kremsmünster, Austria) was used for the large-scale culture. 40 ml of HBSS (-) was used for the final replacement solution according to the ‘**Nutrition-Free Method**’ mentioned in sections **2-1**.

#### 3-2.. Extraction

The collected HBSS (-) after the *“****terminal death****”* of HRC23 cells (**Nutrition-Free Method**) was employed as the resource sample. The sample was centrifuged at 16,000 × g for 10 min (model Avant™ HP-25, 10A rotor, Beckman Coulter™, Inc., CS, USA) and the supernatant was filtrated with a 0.2-µm membrane filter. The filtrate (resource sample) was then immediately dried using a rotary evaporator under a reduced pressure condition [dry diaphragm vacuum pump: KNF Laboport N820.3FT.18(EX) and N810.3FT.18(EX), KNF Neuberger GmbH.] A 1/10 volume of EtOH relative to the resource sample was added to the dried residue, and the supernatant obtained by centrifugation at 1,000 × g for 10 min (KOKUSAN H-108N), was dried again in the same manner. This EtOH extraction was repeated three times to obtain a 1,000-fold concentration of the resource sample. The final EtOH extract was dried in a cryogenic tube and stored at -80 °C for further purification.

#### 3-3.. Column chromatography

Column chromatography was performed under an ambient condition using an HPLC system composed with 880PU pump (×2), 825UV/VIS detector (JASCO Corporation, Tokyo, Japan,) 50 µl through 250 µl Low Flow Series Binary Tee Stainless Steel Housing equipped with 150 μl cartridge, stainless steel (Analytical Scientific Instruments US, Inc., Richmond, CA, ESA,) Rheodyne® 8125 injector (Rheodyne®, Illinois, USA,) and a DG-1310 degasser (Tokyo Rikakikai Co., LTD, Tokyo, Japan.)

##### 3-3-1.. Gel filtration column chromatography

Gel filtration media HP-Cellulofine super fine (sf) [JNC Corporation (Chisso Corporation,) Tokyo, Japan] was packed in a column (Superformance, Ф26mm×L600mm, E·Merck, Darmstadt, Germany.) 50 mM Na_2_SO_4_ was employed as the mobile phase at a flow rate of 0.6 ml/min. For initial gel filtration chromatography, the dried extracted sample was dissolved in 300 μl of the mobile phase and applied to the column via the injector. The elution profile was monitored by absorbance at 230 nm and recorded on a pen-type chart recorder. The elution was collected at 3 min/tube using a fraction collector (Model Frac™ 100, Amersham Biosciences AB, Uppsala, Sweden) and the fractions were divided into eight groups (<u>A</u> to <u>H</u>) on collection. The same volume of MeOH or EtOH was added to the cytocidal active group and Na_2_SO_4_ was removed by passing through a 0.2-µm membrane filter (Millex® GV, Millipore); the sample was dried and stored below -80 °C.

##### 3-3-2.. Ion exchange column chromatography

Two Resource™ S cartridge columns (a strong cation exchange column, GE Healthcare UK Limited, Buckinghamshire, England) were connected in series for use. The gel filtration-processed dried sample was suspended in 300 µl of 0.15 M Na_2_SO_4_ containing 0.01% trifluoroacetic acid (TFA.) The suspension was then applied to the column, followed by washing with 0.15 M Na_2_SO_4_ with 0.01% TFA at a flow rate of 0.2 ml/min for 30 min, and eluted using a linear gradient to 0.24 M Na_2_SO_4_ with 0.01% TFA over 120 min. The elution was monitored by optical absorbance at 230 nm and recorded on the pen-type chart recorder. The elution was collected in 3-minute intervals using the fraction collector. An aliquot of the fraction was withdrawn for cytocidal effect assays, and the cytocidal-exhibited fractions were pooled, dried, and stored at -80 °C. Removal of Na_2_SO_4_ was performed in the same manner as the first gel filtration chromatography.

##### 3-3-3.. Second gel filtration column chromatography

The sample obtained by the ion exchange column chromatography was processed in the same manner as the first gel filtration column chromatography. The elution profile was monitored by absorbance at 205 nm recording on the pen-type recorder. The fraction size was 2 minutes.

### 4.. TOF-MS analysis

TOF-MS analysis was performed using a Voyager DE Pro (Applied Biosystems, CA USA). α-cyano-4-hydroxycinnamic acid [α-CHCA, Tokyo Chemical Industry Co., Ltd., Tokyo, Japan] was used as a matrix. The purified samples, prepared by 2nd gel filtration, were dissolved in 10 µl of sample buffer constituted with 5 mg/ml of α-CHCA in a buffer (acetonitrile: water: TFA (50:50:0.1). Then the sample solutions (0.5 μl) were transferred onto a MALDI plate and dried. The analysis was performed by positive polarity for the sample plate with accelerating voltage at 2000 V and the mass range was from m/z100 to m/z1500 with a bandwidth of 500 MHz.

### 5.. Anti-cancer effect of CSS *in vivo*. Administration to LLC-transplanted mice

Mice (C57BL/6NCrSIc, male, 5 weeks of age) were purchased from Japan SLC, Inc. (Shizuoka, Japan) via the Sankyo Labo Service Corporation, Inc. (Tokyo, Japan). LLC cells (2 × 10^6^ cells), in a 300 µl suspension, were transplanted intraperitoneally using a 30-gauge ×10 mm needle mounted on a 0.5-ml syringe (Nipro Corporation, Osaka, Japan) into four mice (at 6 weeks of age), by our developed method. One week after LLC transplantation, gel filtration processed “CSS” sample, equivalent to 1000ml of the resource sample solution and dissolved in 300 µl of a solution, was injected intraperitoneally into the two mice each, once a day, for six days with free feeding. Another 2 mice were used as non-CSS treated control. Those were followed by continuous observation.

### 6.. Photographs and Microscope

Cells in culture were observed using an inverted microscope model Olympus-CK2 with a 10× objective and an eye lens (Olympus Corporation, Tokyo, Japan). Photographs were taken with a COOLPIX-P310 (NIKON, Tokyo, Japan). Microscopic photos were obtained using an attachment for the camera in place of the eye lens. The optical focal distance was used at 17.9 mm of a zoom range of 4.3 to 17.9mm (28 to 100 mm as 35mm). Scale bar in microscopic photographs was introduced from images of the hemocytometer (improved Burker-Turk) taken in the same manner for the cell’s photographs.

### 7.. Chemicals

All chemicals used were for HPLC quality or higher quality, and cell culture grade throughout this study.

## Key resources table

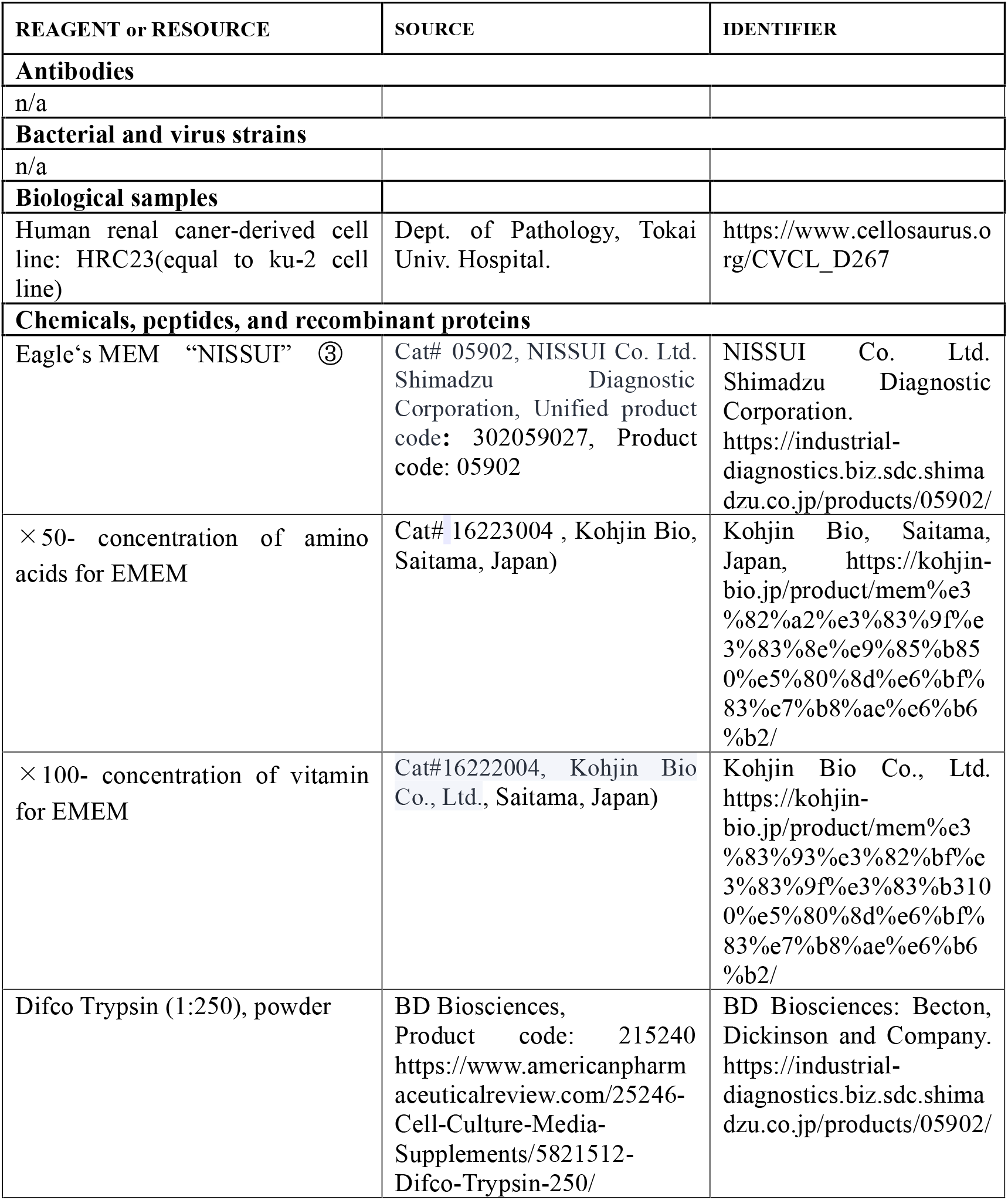
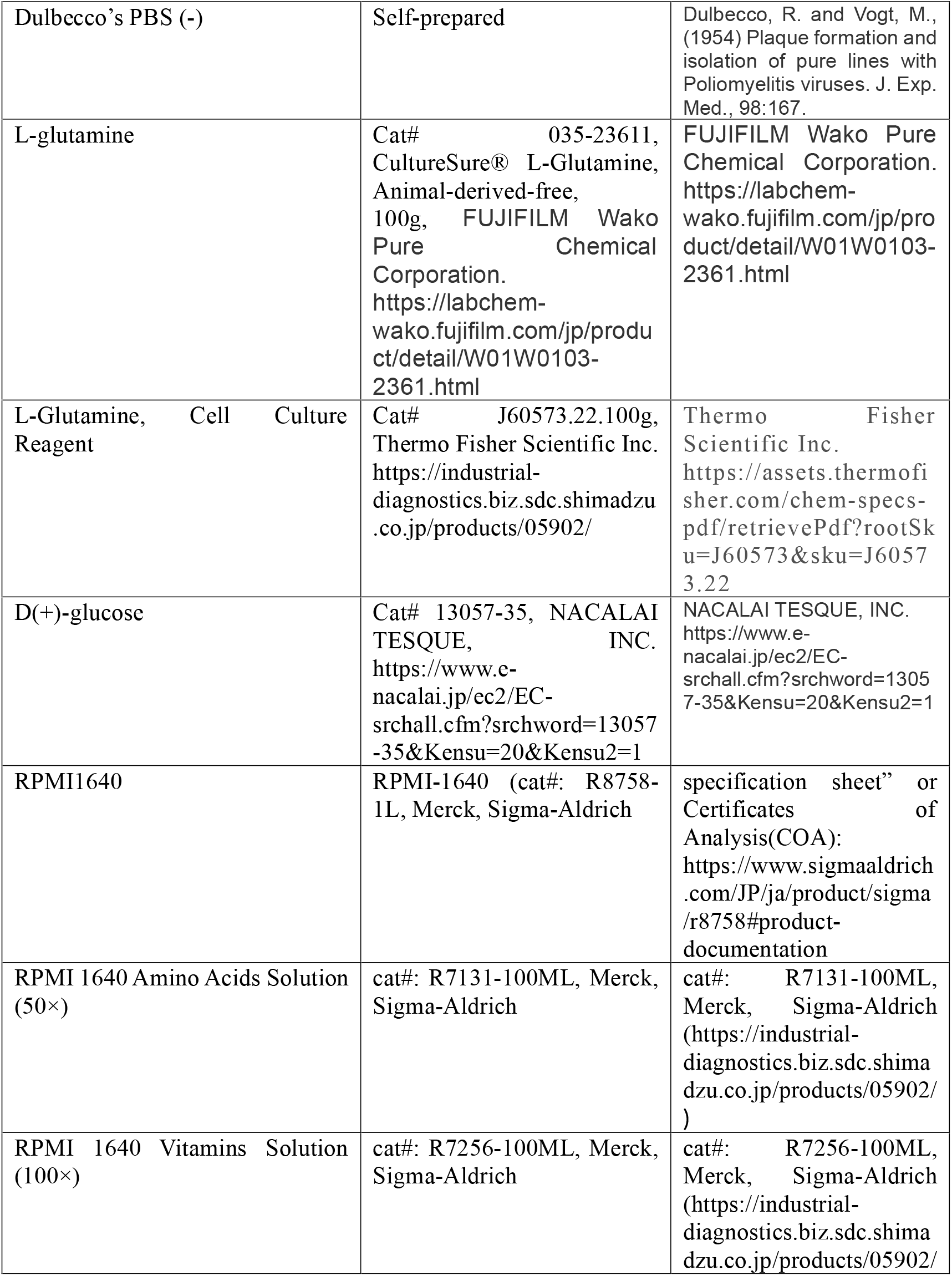
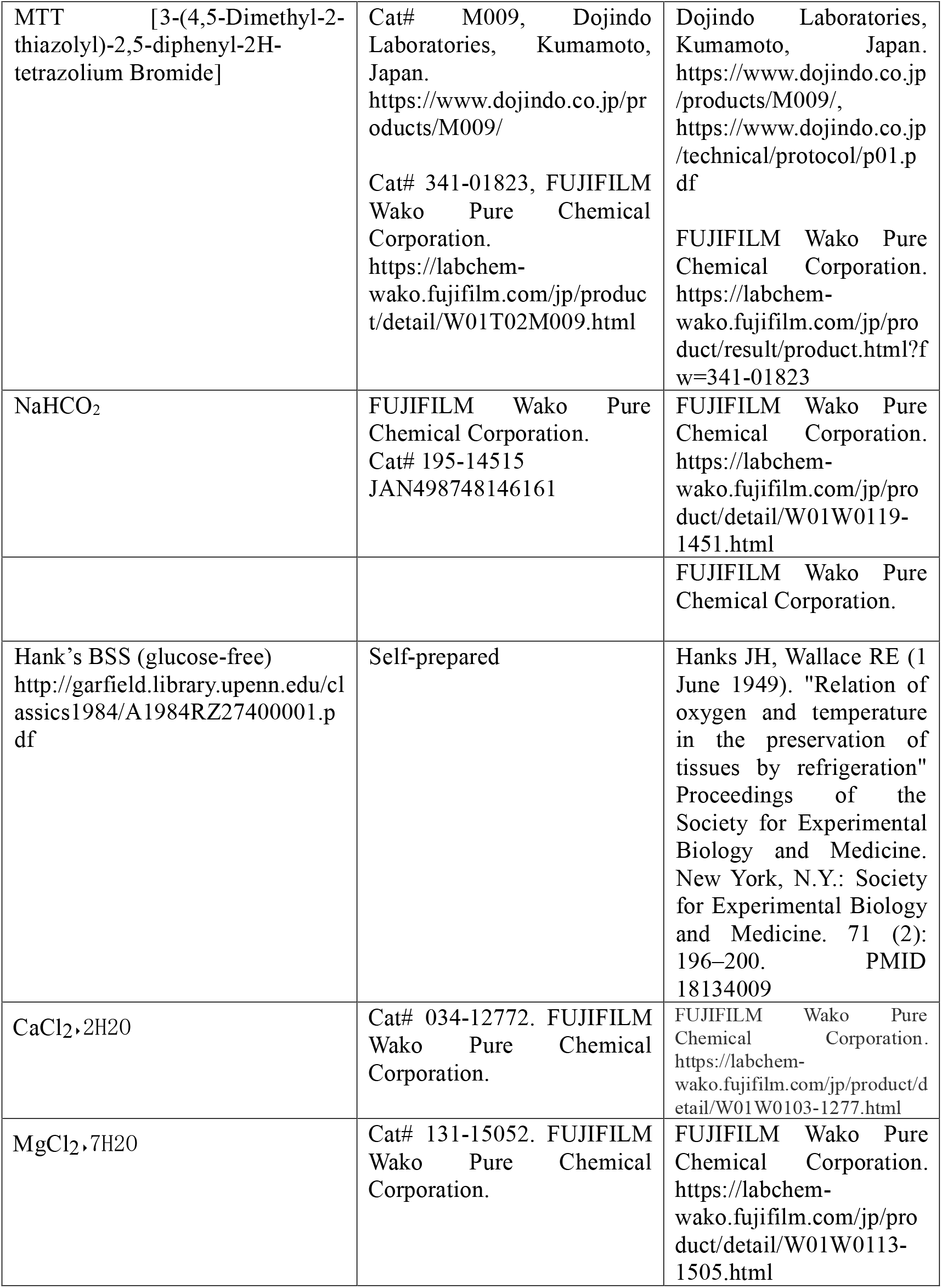
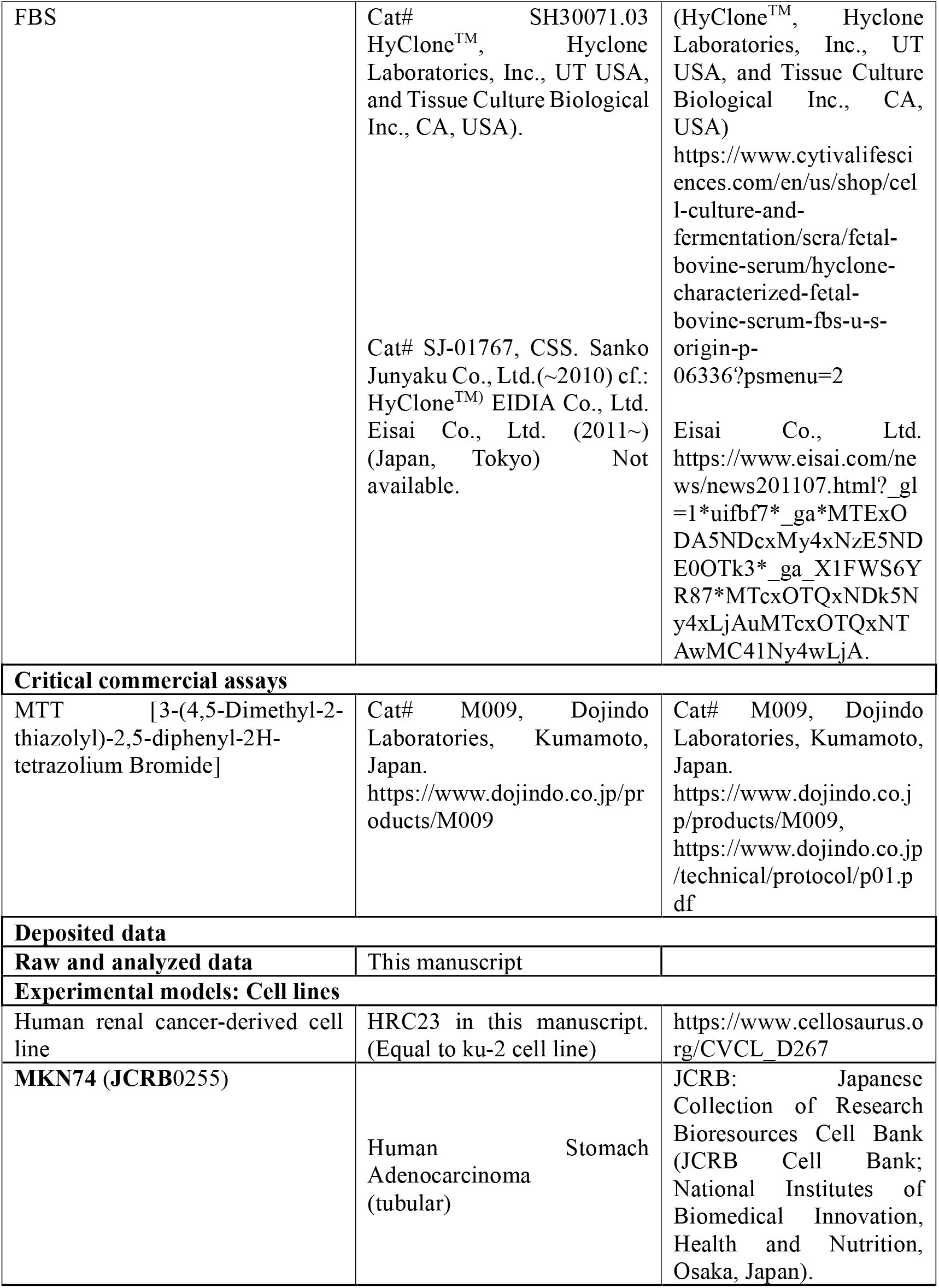
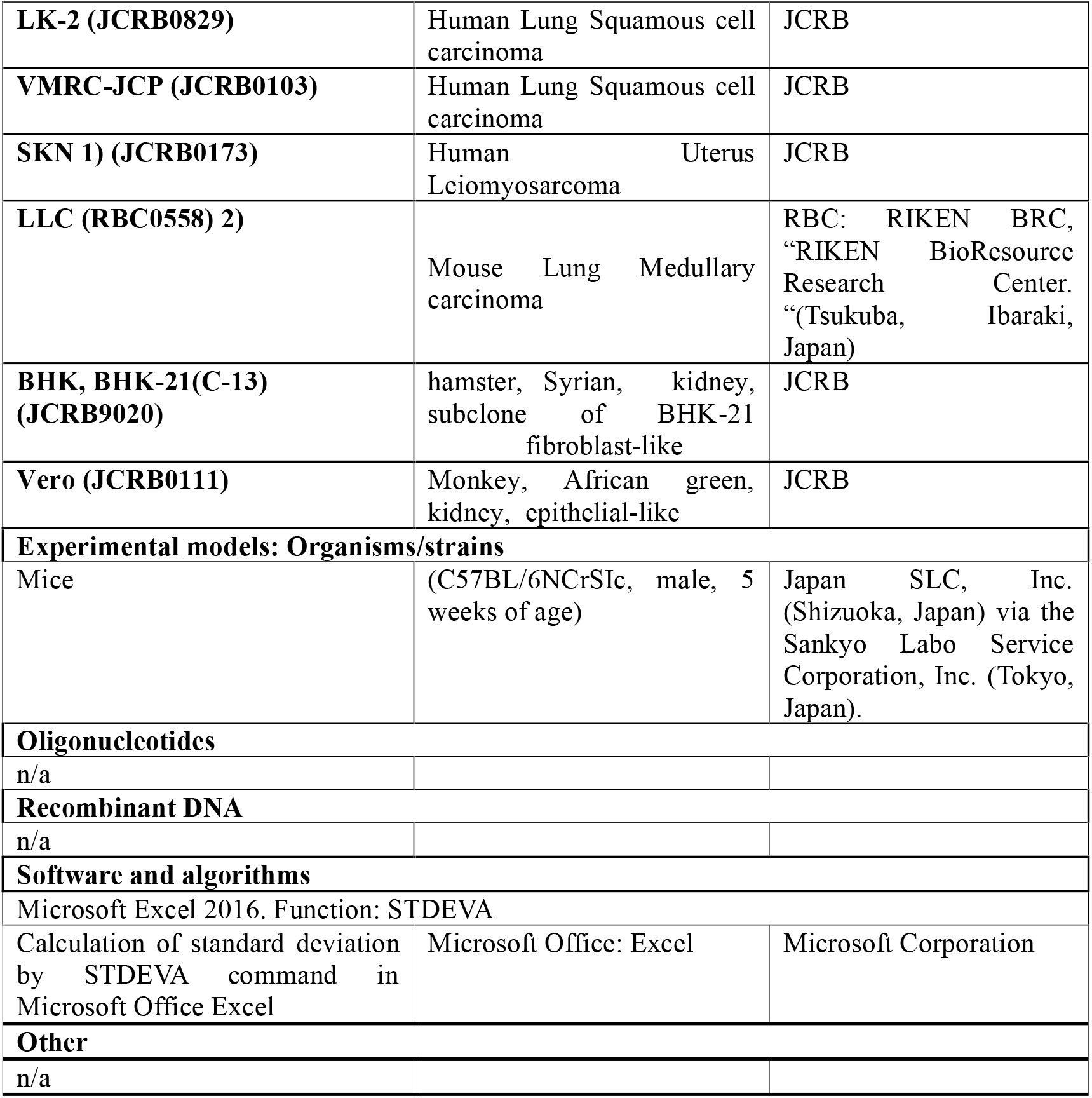

## Notes

**Disclosure of Potential Conflicts of Interest:** The authors declare no potential conflicts of interest. This study was managed only with the income of Medical Corporation Ichikawa Clinic in compliance with Article 3 of the corporation.

### Competing Interest Statement

The authors have declared no competing interest.

